# CytoVI: Deep generative modeling of antibody-based single cell technologies

**DOI:** 10.1101/2025.09.07.674699

**Authors:** Florian Ingelfinger, Nathan Levy, Can Ergen, Artemy Bakulin, Alexander Becker, Pierre Boyeau, Martin Kim, Diana Ditz, Jan Dirks, Jonas Maaskola, Tobias Wertheimer, Robert Zeiser, Corinne C. Widmer, Ido Amit, Nir Yosef

## Abstract

Due to their robustness, dynamic range and scalability, antibody-based single cell technologies, such as flow cytometry, mass cytometry and CITE-seq, have become an irreplaceable part of routine clinics and a powerful tool for basic research. However, their analysis is complicated by measurement noise and bias, differences between batches, technology platforms, and restricted antibody panels. This results in a limited capacity to accumulate knowledge across technologies, studies, experimental batches, or across different antibody panels. Here, we present CytoVI - a probabilistic generative model designed to address these challenges and enable statistically rigorous and integrative analysis for antibody-based single cell technologies. We show that CytoVI outperforms existing computational methods and effectively handles a variety of integration scenarios. CytoVI enables key functionalities such as generating informative cell embeddings, imputing missing measurements, differential protein expression testing, and automated annotation of cells. We applied CytoVI to generate an integrated B cell maturation atlas across 350 proteins from a set of smaller antibody panels measured by conventional mass cytometry, and identified proteins associated with immunoglobulin class-switching in healthy humans. Using a cohort of B cell non-Hodgkin lymphoma patients measured by flow cytometry, CytoVI uncovered T cell states that are associated with disease. Finally, we show that CytoVI is a robust probabilistic framework for the analysis of standard diagnostic flow cytometry antibody panels, enabling the automated detection of tumor populations and diagnoses of incoming patient samples. CytoVI facilitates accurate and automated analysis in both preclinical and clinical settings and is available as open-source software at scvi-tools.org.

## Introduction

Single-cell multi-omics have been applied extensively to assemble comprehensive atlases of individual cells within our body [1], unveil molecular and cellular perturbations that may lead to the manifestation of disease or explain a patient’s response to therapy [2]. Having emerged as an irreplaceable component of biomedical research, these technologies utilize the cells’ transcriptome, epigenome, proteome or morphology as a proxy to determine their cells intrinsic state that may ultimately translate into function. Pre-dating the explosion in use of large-scale genome-wide protocols, flow cytometry was the first technology capable of analyzing millions of individual cells within minutes [3]. Due to its low cost, dynamic range, robustness and scalability, flow cytometry has become an irreplaceable part of clinical routine, facilitating the diagnosis of hematological malignancies or serving as a prognostic tool to guide treatment decisions.

Since its introduction in 1965, flow cytometry has dramatically advanced. The increase of more than 50 cellular features that can be simultaneously analyzed and its cost effectiveness have transformed flow cytometry to a powerful tool for exploratory research [4]. However, increasing the dimensionality of flow cytometry faces inherent technical challenges, primarily due to the spectral overlap of fluorophores. As the number of detectable markers grows, distinguishing between fluorochromes with overlapping emission spectra becomes increasingly complex. This limitation not only necessitates complex unmixing strategies but also imposes a fundamental physical constraint on the further development of fluorescence-based cytometry platforms. The application of heavy-metal tagged antibodies in mass cytometry (CyTOF) or oligonucleotide-tagged antibodies using Cellular Indexing of Transcriptomes and Epitopes by sequencing (CITE-seq) held great promise to further increase the number of proteins that can be detected per cell [5, 6]. However, challenges in labeling and chelator chemistry have limited the dimensionality of mass cytometry assays. Economic limitations of oligonucleotide-tagged antibody assays followed by sequencing have prevented CITE-seq from being utilized in a similar scale to measure protein abundance in a comparable throughput and sensitivity as for flow and mass cytometry. Consequently, the expansion of antibody-based single-cell assays are currently constrained by a combination of physical, chemical, and economic factors, unique to each approach.

The combination of machine-learning and traditional flow cytometry demonstrated the tremendous potential to computationally mitigate the limited number of cellular features detectable, but was not practicable for multi-donor settings due to the requirement of a high number of cells per assay [7]. Additionally, deep generative latent variable models have emerged as powerful tools for the integration, denoising, and imputation of single-cell genomics data [8]. These models have significantly expanded the scope of biological questions that can be addressed by enabling more accurate inference of cellular states, trajectories, and regulatory networks from inherently sparse and noisy data [9]. In contrast to the stochastic nature and sparsity commonly associated with genomic single-cell technologies, antibody-based cytometry technologies offer high-resolution measurements of millions of cells across a wide dynamic range facilitating the analysis of large patient cohorts. However, the analysis of multi-cohort studies is often obstructed by batch effects attributed to differences in sample isolation protocols or antibody staining conditions. Moreover, comparative studies using the vast amount of publicly available cytometry data are hampered by different technologies applied to generate data and the use of distinct antibody panels utilized to interrogate samples.

Here, we present CytoVI, a probabilistic end-to-end deep learning framework designed for the integration of antibody-based single cell technologies. CytoVI removes technical variation in flow cytometry, mass cytometry or CITE-seq data and embeds cells into a meaningful low-dimensional representation corresponding to a cell’s intrinsic state. We demonstrate that CytoVI is capable of imputing missing markers in experiments with overlapping antibody panels, impute a cell’s transcriptome if paired with CITE-seq data and facilitate the automated detection of cellular states that are associated with clinical covariates, such as a patient’s response to treatment. Thus, CytoVI enables the integration of the different methodiolgies, taking advantage of the throughput and scalability of cytometry approaches with the dimensioanlity of genome wide technologies. CytoVI is available as an open-source Python library at scvi-tools.org [10], leveraging advanced optimization strategies, including multi-GPU training and efficient data-loading mechanisms that enable direct access to a large collection of datasets. These features facilitate the scalable modeling of hundreds of millions of cells. CytoVI mitigates some of the most pronounced limitations associated with modern cytometry and single cell genomics technologies and paves the way towards a next generation of AI-powered cytometry integrated with other single cell genomic modalities with tremendous potential for biomarker discovery and applicability in clinical practice.

## Results

### The CytoVI model

Flow cytometry is arguably the most abundant single cell data modality in the public domain and thus has a great potential for cellular atlassing of many thousands of individuals across billions of cells. However, efficient large-scale integration of antibody-based single cell technologies is hampered by technical variation introduced during data generation. To decouple technical variation from biological variation in a unified manner for antibody-based single cell technologies, CytoVI encodes each cell’s protein expression in a shared normally-distributed latent space that corresponds to its intrinsic state with its associated uncertainty **(Figure 1A)**. In a subsequent step, CytoVI generates probabilistic estimates of the protein expression profile of each cell, given its embedding coordinates. In addition to the protein expression matrix, CytoVI can incorporate observed nuisance factors that inform of technical variation due to experimental batches, differences in the applied technology, or confounding biological variables such as sex or age. These artifacts are controlled for in the inferred embedding and in the generative process, thus giving rise to corrected estimates of cell state and protein expression. Another optional input for CytoVI are cell type labels, computed *a priori*. CytoVI uses these labels to inform the prior distribution of the embedding space (with different modes associated with different cell types; see Methods), a step which we generally found helpful in difficult integration scenarios [11].

**Figure 1.**
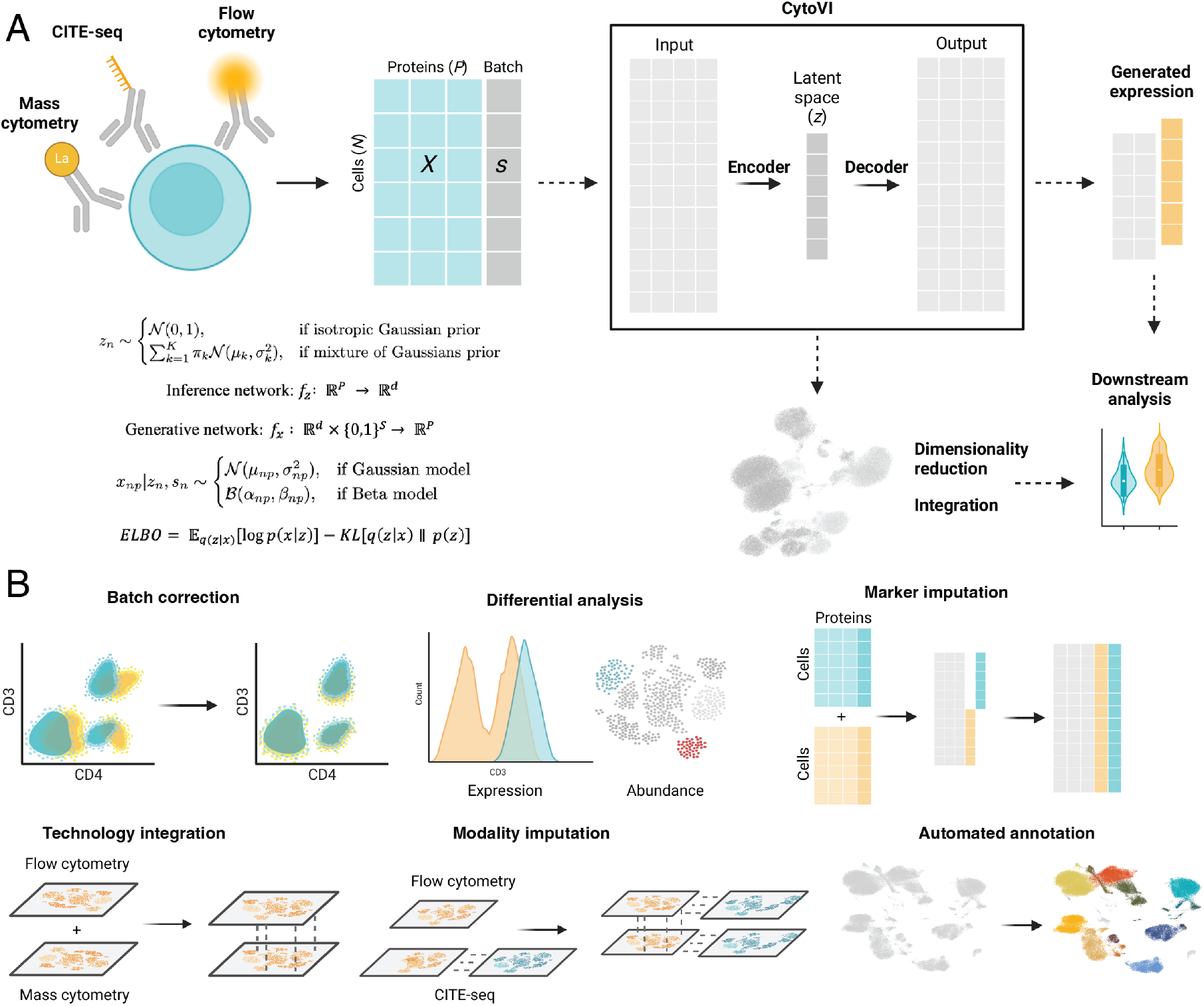
Schematic overview of CytoVI. **A:** CytoVI is a latent variable model consisting of an encoder and a decoder neural network. The encoder maps the protein expression matrix of antibody-based single cell technologies (*X*) into a joint latent space (*z*). Depending on the expected underlying biology of the data we assume an isotropic Gaussian prior or a mixture of Gaussians prior for the posterior distribution *p*(*z*|*x*). The decoder network incorporates the posterior mean of (*z*) and observed sample-level covariates such as batch (*s*) and fits the parameters of a Beta- or Gaussian distribution for the protein likelihood in order to reconstruct the original protein expression. The variational and neural network parameters are trained by maximizing the *ELBO* consisting of a reconstruction term and Kullback-Leibler (KL) divergence. Thereby, CytoVI optimizes a latent representation for each cell that is corrected for the observed sample-level covariates (*s*). **B:** Schematic overview of the analysis tasks that can be accomplished using CytoVI.

Both the encoder that facilitates inference of latent distributions and the decoder, corresponding to the generative model that estimates the distribution of a cell’s protein expression, are comprised of neural networks. Amortized variational inference estimates the neural network parameters and the variational parameters of the distribution in latent space and protein expression likelihood. After training, the CytoVI latent space can be utilized for data integration, cell type assignment or label-free differential abundance analysis. Conversely, the generative model can be utilized to obtain corrected expression estimates, identify differentially expressed proteins or impute missing markers when overlapping antibody panels are applied **(Figure 1B)**. For the latter cases in which data generated using distinct antibody panels are integrated, the inference network utilizes only the shared overlapping markers to optimize a cell’s latent distribution but the generative network reconstructs the union of all markers present in the antibody panels, thereby effectively imputing unseen markers (which are masked during computation of the reconstruction loss).

The signal magnitude of cytometry data heavily depends on the choice of reporter molecules tagged to the respective antibody (fluorophore or heavy metal isotope). In contrast to most single cell genomics modalities, it does not provide absolute measurements of protein abundance but a relative quantification in relation to other cells within the assay. In addition to the indirect quantification of relative protein abundance using antibodies, flow cytometry captures scatter features that constitute label-free surrogate markers for cell size and granularity. To account for the relative nature of antibody-based single cell data and to improve its suitability for visualization and downstream analysis (particularly by making the transformed data more comparable to a Gaussian distribution), cytometry data is commonly preprocessed using hyperbolic arcsin, logicle, or biexponential transformations [12, 13]. These transformations are followed by scaling to normalize the magnitude of individual marker expression to a comparable range.

Given this preprocessing approach, we reasoned that the noise model of a cell’s protein expression after hyperbolic arcsin transformation, e.g. the distribution when repetitively sampling the protein expression of the same cell, could be approximated using a Gaussian distribution. To empirically test this assumption, we measured the fluorescence intensities of homogeneous microspheres that either captured a uniform number of fluorescent antibodies or did not bind any antibodies, serving as negative controls **(Figure S1A)**. Indeed, after hyperbolic arcsin transformation, fluorescence intensities of antibody-labeled microspheres displayed bimodal peaks that could be approximated using a Gaussian distribution **(Figure S1A and B)**. To confirm that hyperbolic arcsin transformed antibody-based single cell data across technologies could be represented as a Gaussian function of a cell’s state, we next assessed the distribution of frequently assessed lineage and activation markers in peripheral blood mononuclear cells (PBMCs) of healthy donors measured by flow cytometry, mass cytometry and CITE-seq [14, 15, 16]. After transformation, the distribution of marker intensities were comparable between different technologies and could be approximated using a mixture of Gaussian distributions (with modes distinguishing between cell states/types), supporting the applicability of a joint latent variable model between the different technologies **(Figure S1C and D)**. Accordingly, the generative model of CytoVI relies on Gaussian distribution. We also allow for beta distribution if the transformed expression values are scaled between zero and one.

### CytoVI accurately models antibody-based single cell data and provides a biologically interpretable latent space

To evaluate the ability of CytoVI to generate data that closely fit the observed cytometry data, we compared CytoVI using different protein likelihood and preprocsessing options to factor analysis, a linear Gaussian baseline method proposed by our earlier work [17]. We performed posterior predictive checks (PPCs) and measured how well the coefficient of variation (CV) of the generated data corresponded to the observed CV in the flow cytometry dataset of healthy PBMCs [14]. Each of the variants of CytoVI outperformed the factor analysis in its ability to fit the observed flow cytometry data **(Figure S2A)**, with the configuration utilizing a Gaussian likelihood and operating on hyperbolic arcsin transformed and scaled data (as recommended as best practice) [18, 12] showing the best performance **(Figure S2A)**. To assess whether the generative model of CytoVI could generalize to antibody-based single cell technologies beyond flow cytometry, potentially enabling integration with data of higher dimensionality, we next performed PPCs of the different CytoVI configurations and the factor analysis baseline for PBMCs meaured by mass cytometry and oligonucleotide-tagged antibody data (CITE-seq) [16, 15]. CytoVI generated protein expression data that closely fitted the observed data. The configuration utilizing the Gaussian likelihood best preserved the mean-variance relationship of the data and was thus selected as default configuration **(Figure S2B and C)**. Collectively, we demonstrated CytoVI’s ability to generate protein expression data that closely aligned with observed measurements from flow cytometry, mass cytometry and CITE-seq data.

Next, we evaluated the biological interpretability of CytoVI’s latent space and the extent to which it captures cell state variability across the different technologies. For this, we applied clustering to the protein expression (unintegrated data space) and used key protein markers to manually assign cell type labels for the human PBMC datasets measured using flow cytometry, mass cytometry and CITE-seq [14, 15, 16] **(Figure S3A and B)**. In case cell type labels have been provided with the original study, these cell type labels have been retained **(Figure S3C)**[16]. Visualizing these cell type annotations in the CytoVI latent space confirmed its applicability to resolve cell type variation across flow cytometry, mass cytometry and CITE-seq data **(Figure 2A)**. Next, we evaluated whether the CytoVI latent space could be employed for *de novo* cell type annotation. For this, we performed Leiden clustering on the CytoVI latent space and assessed the extent to which these clusters recapitulate the manually-assigned cell type annotations. Clusters obtained from the CytoVI latent space largely replicated the prior cell type annotations of even rare subsets, such as dendritic cells (*<*2%), while avoiding over-fragmentation of larger populations as observed in other computational workflows for cytometry analyses **(Figure 2B, Figure S3D and E)** [19, 20]. Moreover, the CytoVI latent space comprised fine-grained cell state variability beyond the manual annotation level as exemplified by protein expression gradients in the natural killer (NK) cell compartment. CD56^dim^ NK cells exhibited a gradient of CD16 expression, the IgG receptor mediating antibody-dependent cytotoxicity, which closely paralleled the expression of the inhibitory receptor TIM-3. In contrast, an opposing expression pattern was observed for the cytotoxic molecule Granzyme B **(Figure 2C)**. These observations align with previous reports demonstrating that TIM-3 is upregulated following Fc-receptor engagement of CD16, thereby impairing the cytotoxic capacity of NK cells [21, 22]. Together, we demonstrate that CytoVI optimizes a biologically interpretable latent space, capturing a cell-intrinsic state that can be utilized for cell type annotation or for identification of gradual protein expression changes.

**Figure 2.**
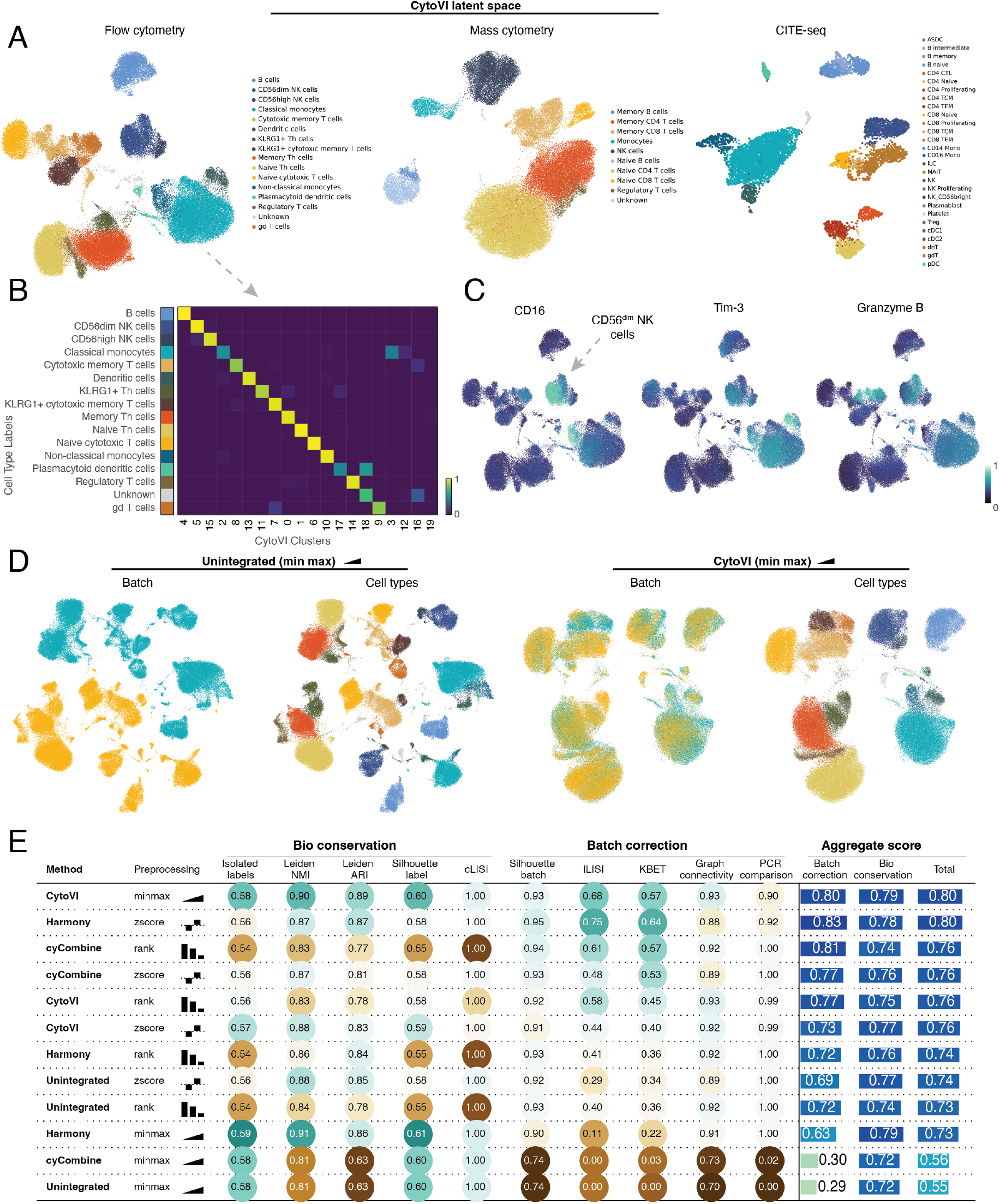
CytoVI optimizes a biologically-interpretable latent representation for antibody-based single-cell technologies that controls for technical variation. **A:** UMAP of the CytoVI latent space for PBMCs measured by flow cytometry (left panel), mass cytometry (middle panel), and CITE-seq (right panel). Cells are colored by the ground truth cell type annotations that have been either provided with the original publication of the data (right panel) or were manually annotated. **B:** Heatmap displaying the confusion matrix between the ground truth cell type annotations of the flow cytometry dataset and clusters obtained by clustering the CytoVI latent space. **C:** UMAP of the CytoVI latent space of the flow cytometry dataset indicating the expression for CD16, Tim-3, and Granzyme B. **D:** UMAP of the unintegrated data space of two biological replicates of flow cytometry samples measured in two different batches, colored by batch and the respective cell type annotation (left panels). UMAP of the CytoVI latent space of the same data after controlling for the observed batch covariate (right panels). In both cases, the arcsinh-transformed data has been min-max scaled to ensure equal representation of each marker. **E:** Overview table benchmarking the integration performance of CytoVI, cyCombine, and Harmony for the flow cytometry dataset in relation to different preprocessing choices. Different metrics are presented for bio conservation and batch correction, and an aggregated consensus score summarizes the results.

### CytoVI removes technical variation while preserving biological variation

Batch effects, arising from differences in experimental protocols, staining conditions, instrument performance or reagents lots, are a major challenge in flow cytometry analyses [12]. Therefore, we next assessed the extent to which CytoVI can control for these observed nuisance factors, while simultaneously preserving biological variation. For this, we utilized replicates of cryopreserved PBMCs that have been analyzed in two distinct batches by flow cytometry and demonstrated a clear batch effect [14]. When explicitly controlling for the observed batch covariate, CytoVI yielded a latent space that integrated both batches while preserving cell type variation **(Figure 2D)**. Next, we benchmarked CytoVI’s integration performance against cyCombine, a recently proposed method for the integration and imputation of cytometry data, and Harmony, a commonly applied integration method introduced for single cell genomics that has been heavily applied for the integration of cytometry data [23, 24]. cyCombine required the cytometry data to be standardized first (e.g. using z-scores). For comprehensive evaluation we therefore conducted our tests with three different standardization methods **(Figure 2E)**.

We evaluated different metrics for batch correction and conservation of biological information for each integration methodology and standardization method using the single cell integration bench-mark framework [25]. We find that standardization with z-scores or ranks, with no additional modeling, was able to reduce the technical variation to some extent, while min-max transformation retained the batch effects (**Figure 2E** and **Figure S3F**). Adding the integration component, we observed that cyCombine and Harmony improved the integration of Z-scored or rank-transformed data, but failed to do that with the min-max transformation. Conversely, CytoVI led to a clear improvement in all three standardization approaches and generally compared favorably to the other methods. Taking a closer look (beyond the summary scores in **Figure 2E**) we find that cyCombine could not account for the technical variation in the myeloid compartment, while Harmony maintained an apparent batch effect in the T cell compartment **(Figure S3F)**. These specific nuisance effects are not observed with CytoVI. Together, we demonstrate that CytoVI excels at integration tasks and outperforms current state-of-the-art integration tools for cytometry data.

### CytoVI facilitates accurate imputation of unobserved proteins

The combination of traditional flow cytometry and machine-learning has demonstrated marked potential to overcome the limitation of the low dimensionality of cytometry assays [7]. CytoVI’s generative model comprises an intrinsic ability to generate unobserved data and thus to impute missing proteins from cytometry experiments, where different antibody panels have been applied **(see Methods)**. To evaluate CytoVI’s performance for imputing unobserved proteins, we generated semi-synthetic batches by partitioning flow cytometry data into two identical batches and masked one marker at the time in one of the batches. Consecutively, the missing marker was imputed using CytoVI’s generative network and compared to the held-out observed protein expression in each cell. This procedure was iterated for each marker and CytoVIs imputation performance was compared to KNN imputation (in data space) and cyCombine’s imputation module [23, 26]. Of note, the overall imputation performance varied between individual markers. Lineage markers such as CD4, CD14 or CD3 were imputed with high accuracy (Pearson’s r *>* 0.9), while Ki67 or CD103 achieved lower imputation results (Pearson’s r *>* 0.6) **(Figure S4A and B)**. Yet, across all markers analyzed CytoVIs imputation performance was superior to cyCombine **(Figure S4B)**. Similar benchmarking results were obtained when evaluating the ability to impute the binary marker positivity as often practiced in flow cytometry data analysis **(Figure S4C and D)**. The variable results associated with the imputation of markers and the corresponding uncertainty about accuracy of imputed values have limited the application of protein imputations in the field of cytometry to specific cases so far.

However, due to its probabilistic nature, CytoVI provides an intrinsic estimate about imputation uncertainty that strongly correlated with the *de facto* imputation error **(Figure S4E)**. Therefore, CytoVI not only provides state-of-the-art imputations for missing protein markers in overlapping antibody panels but also estimates the uncertainty of imputations, which can be utilized to assess reliability of protein predictions for downstream analyses.

To demonstrate CytoVI’s applicability to integrate data from distinct experiments comprising different antibody panels, we integrated the flow cytometry PBMC dataset from *Nunez et al*. with additional flow cytometry data of PBMCs obtained from *Kreutmair et al* [27] **(Figure 3A)**. To incorporate our prior knowledge about cell types present in both datasets, we included the cell type labels to inform the prior of the CytoVI latent space (**Methods**). The CytoVI latent space effectively controlled for the technical variability between the two experiments but retained donor-specific cell states, such as the abundance of *γδ* T cells and plasmablasts that can be highly variable between individuals **(Figure 4B, Figure S5A)** [28, 29]. Imputation of non-overlapping proteins increased the dimensionality of the assay and facilitated a deeper interrogation of immune cell states. This was exemplified by immunophenotyping of CD56^dim^ CD16^+^ NK cells that expressed the immune checkpoint receptor KLRG1 (measured only in the *Nunez et al*. dataset) and the C-type lectin receptor CD161 (measured only in the *Kreutmair et al*. dataset). A smaller subpopulation of CD56^dim^ CD16^+^ NK cells expressed the maturation marker CD57 (measured only in the *Kreutmair et al*. dataset) **(Figure 3B, S5B)**. This is in line with a previous single cell RNA sequencing atlas describing NK cell subsets in the peripheral blood of healthy donors that proposed CD57^+^ NK cells as a distinct molecular subset of mature human NK cells [30]. Merging of the two antibody panels using CytoVI accumulated knowledge about the protein expression from both datasets and enhanced the capability to perform comprehensive immunophenotyping (e.g. by generating large-scale protein atlases). In conclusion, we have demonstrated the integration and imputation of flow cytometry data from two distinct studies, thereby increasing the number of proteins that could be interrogated, and facilitating a shared downstream analysis **(Figure 3D)**.

**Figure 3.**
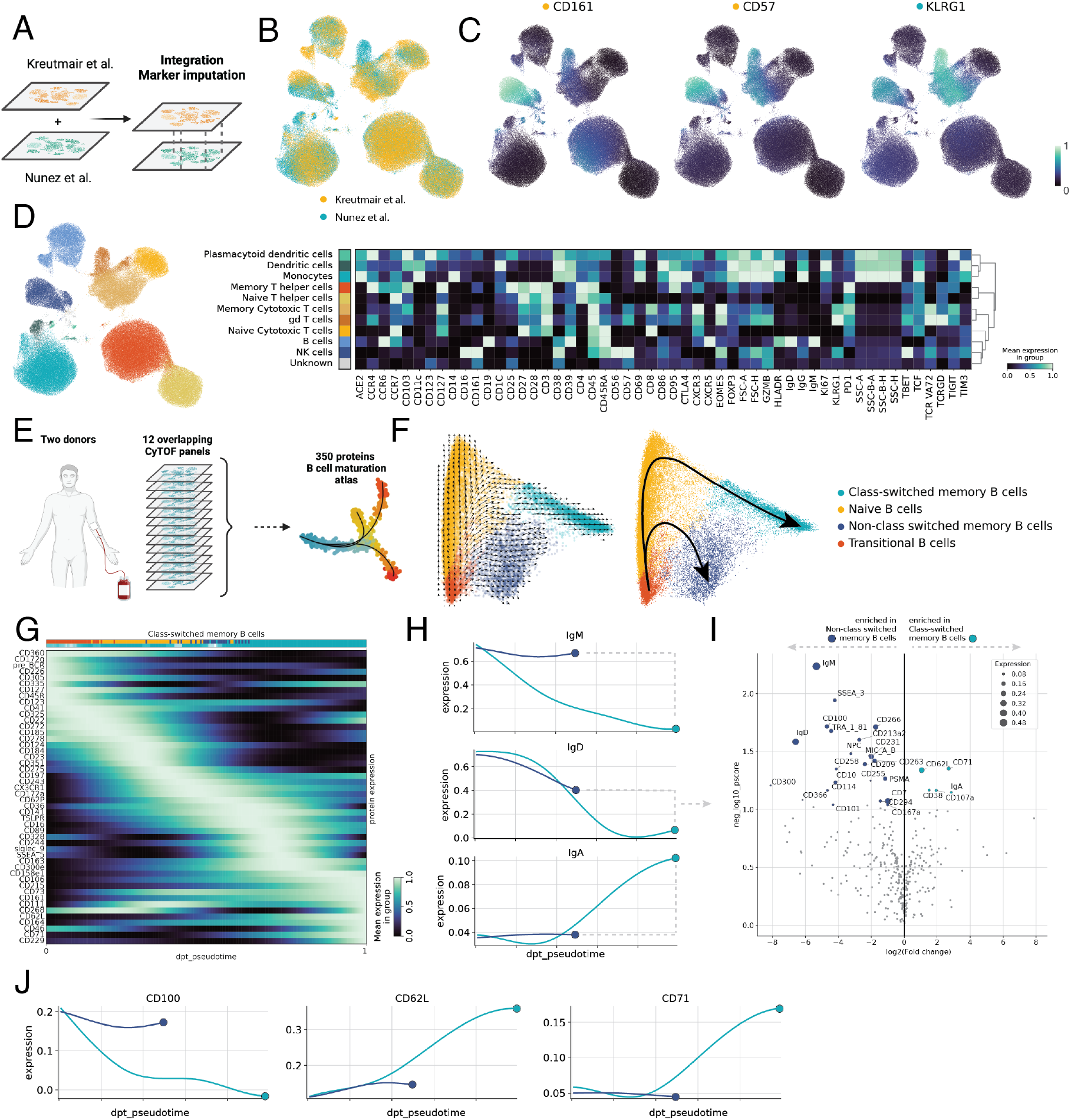
CytoVI integrates cytometry data from distinct antibody panels and imputes missing proteins. **A:** Schematic overview for the integration of two distinct flow cytometry studies from *Nunez et al*. (35 markers) and *Kreutmair et al*. (30 markers) with a shared overlap of 15 backbone proteins markers. **B:** UMAP of the CytoVI latent space colored by the respective study. **C:** UMAP of the CytoVI latent space displaying the imputed expression of CD161, CD57 (both measured in the *Kreutmair et al*. study) and KLRG1 (measured in the *Nunez et al*. study). **D:** UMAP of the integrated CytoVI latent space colored by harmonized cell type annotations between the two studies (left panel). Heatmap showing the imputed protein expression profile for the indicated cell types across the union of proteins measured in both studies. **E:** Schematic overview of the integrated B cell maturation atlas from mass cytometry data generated by *Glass et al*.. **F:** Diffusion maps of the imputed protein expression profile of 350 surface markers showing the cell-cell transition matrix obtained from CellRank (left panel) and a manual summary of the B cell maturation trajectories (right panel). Coloring indicates cell type annotations obtained from clustering the imputed B cell maturation atlas. **G:** Heatmap displaying the imputed protein expression of the top 50 markers that correlate best with the differentiation trajectory from transitional B cells to class-switched memory B cells along the diffusion pseudotime. **H:** Line plots displaying the expression of the indicated immunoglobulins for the class-switched (teal) and non-class-switched memory B cell lineage (blue) along the diffusion pseudotime. **I:** Volcano plot displaying the differentially expressed proteins between class-switched (teal) and non-class-switched (blue) memory B cells determined using CytoVIs differential protein expression module. **J:** Line plots displaying the expression of the indicated proteins for the class-switched (teal) and non-class-switched memory B cell lineage (blue) along the diffusion pseudotime.

**Figure 4.**
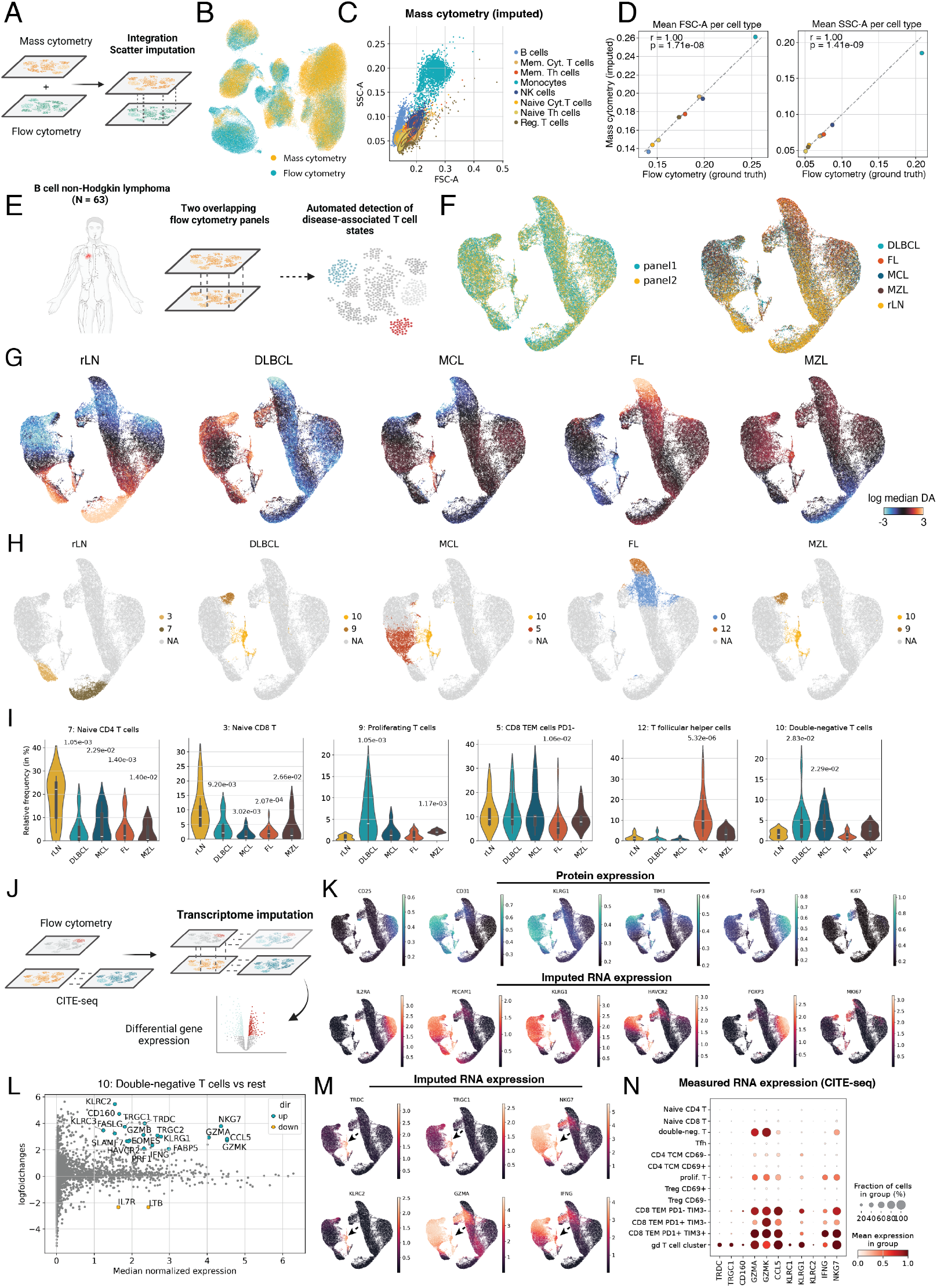
CytoVI facilitates cross-technology integration and imputation of missing modalities. **A:** Schematic overview for the integration of flow and mass cytometry data. **B:** UMAP of the CytoVI latent space displaying the integration of mass (yellow) and flow (teal) cytometry data. **C:** Scatter plot displaying the imputed morphological features FSC and SSC. Cells are colored by the indicated cell types. **D:** Scatter plot showing the mean FSC and SSC for the indicated immune population measured by flow cytometry (ground truth) and imputed in mass cytometry data. Statistics refer to Pearson’s correlation. **E:** Schematic overview of the B cell non-Hodgkin lymphoma analysis. **F:** UMAP of the CytoVI latent space of lymph node samples of B cell non-Hodgkin lymphoma patients colored by the antibody panel utilized to generate the data (left panel) or the indicated lymphoma entities (right panel). rLN = resected lymph node (N = 7), DLBCL = diffuse large B cell lymphoma (N = 17), MCL = Mantel cell lymphoma (N = 9), MZL = marginal-zone lymphoma (N = 6), FL = follicular lymphoma (N = 24). **G:** UMAP of the CytoVI latent space displaying the differential abundance scores for the indicated disease entities. **H:** UMAP of the CytoVI latent space highlighting the top two differentially abundant clusters for each disease entity. **I:** Violin plots displaying the relative frequencies of the indicated differential abundance clusters across the different disease entities. P values refer to pair-wise Mann-Whitney U tests in relation to resected control lymph nodes (rLN). **J:** Schematic overview displaying the integration of the flow cytometry cohort with paired CITE-seq data for imputation of the transcriptome. **K:** UMAP of the CytoVI latent space showing the protein expression of the indicated markers measured by flow cytometry (upper panel) and the imputed gene expression of the corresponding transcripts (bottom panel). **L:** MA plot ranking the log fold change of imputed gene expression of the double-negative T cell cluster compared to all other T cells in relation to the median normalized gene expression of the double-negative T cell cluster. **M:** UMAP of the CytoVI latent space showing the imputed gene expression of the indicated transcripts. **N:** Dot plot displaying the scaled normalized gene expression profile of the *γδ* T cell cluster detected in the CITE-seq dataset in relation to all other T cell subsets. FSC = forward scatter, SSC = side scatter.

### CytoVI assembles an integrated B cell maturation atlas spanning 350 protein markers

Next, we evaluated the extent to which CytoVI could be leveraged to construct large-scale protein atlases from conventional cytometry assays. To this end, we analyzed mass cytometry data of circulating B cells from two donors, measured across 12 partially overlapping antibody panels. Each of the panels provides a limited view, with approximately 39 targeted proteins, but their union results in a comprehensive set of 350 markers [31] **(Figure 3E)**. Five of the nine backbone markers (markers shared between all antibody panels) that could be utilized to define B cell subpopulations exhibited variable distributions across samples, indicating the presence of batch effects **(Figure S5C)**. However, when explicitly controlling for the batch covariate, CytoVI generated a latent space that minimized technical variation, while preserving biological variation across B cell states **(Figure S5D and E)**. Clustering of this latent space and imputation of the non-overlapping protein markers enabled the in-depth characterization of 350 surface proteins across transitional, naïve, non-class-switched, and class-switched memory B cells in the human peripheral blood **(Figure S5F)**. Notably, relative proportions of B cell subsets in the two donors remained unaffected by data integration, suggesting that integration preserved biological variability **(Figure S5G)**.

Since B cell maturation and differentiation follow a continuous trajectory rather than discrete clusters, we next modeled the differentiation route as a dynamic biological process. For this, diffusion maps of the integrated B cell atlas were computed and the cell-cell transition matrix was estimated using CellRank’s pseudotime kernel based on the diffusion pseudotime [32, 33]. Visualization of the cell-cell transition matrix and estimation of initial and terminal states revealed that transitional B cells mature into naïve B cells and consecutively commit either to the non-class switched memory B cell fate (such as marginal zone-like B cells) or differentiate into class-switched memory B cells **(Figure 3F, Figure S5H and I)**. Notably, analysis with Palantir pseudotime yielded comparable results as for diffusion pseudotime [34, 35] **(Figure S5H)**. Next, we queried our B cell maturation atlas and identified driver proteins that best associate with the differentiation towards the class-switched memory B cell fate by correlating the individual protein expression of a cell with its probability to differentiate into the class-switched memory B cell terminal state **(see Methods)**. This revealed protein expression gradients crucial for maturation of B cells. For instance, transitional B cells migrating from the bone-marrow into the circulation displayed rapid downregulation of the pre-B cell receptor, followed by transient decay of CD22, a negative regulator of B cell receptor signaling, and CD185 (CXCR5), which is required for germinal center formation **(Figure 3G)**. Lineage commitment was most apparent when visualizing the expression of immunoglobulins (the hallmark of class-switching) that demonstrated the switch from IgD and IgM towards IgE in the class-switched memory B cells, while non-class switched memory B cells retained surface IgD and IgM **(Figure 3H)**. To identify proteins that are specifically associated with the class-switched memory B cell lineage compared to non-class switched memory B cell fate, we utilized CytoVI’s generative model and tested for differentially expressed proteins. This revealed proteins involved in class-switching such as CD62L, a selectin essential for migration into lymph nodes [36], CD71, a proposed marker for antigen-specific B cells [37] or CD100, which has been shown to modulate CD40-CD40L interactions [38] **(Figure 5I and J)**. These results highlight the role of B cell/T cell interactions required for somatic hypermutation and isotype class-switching in B cells. In conclusion, we demonstrate CytoVI’s performance in assembling large-scale protein atlases from conventional cytometry assays that can be utilized to identify biomarkers or uncover novel regulators of complex biological processes.

**Figure 5.**
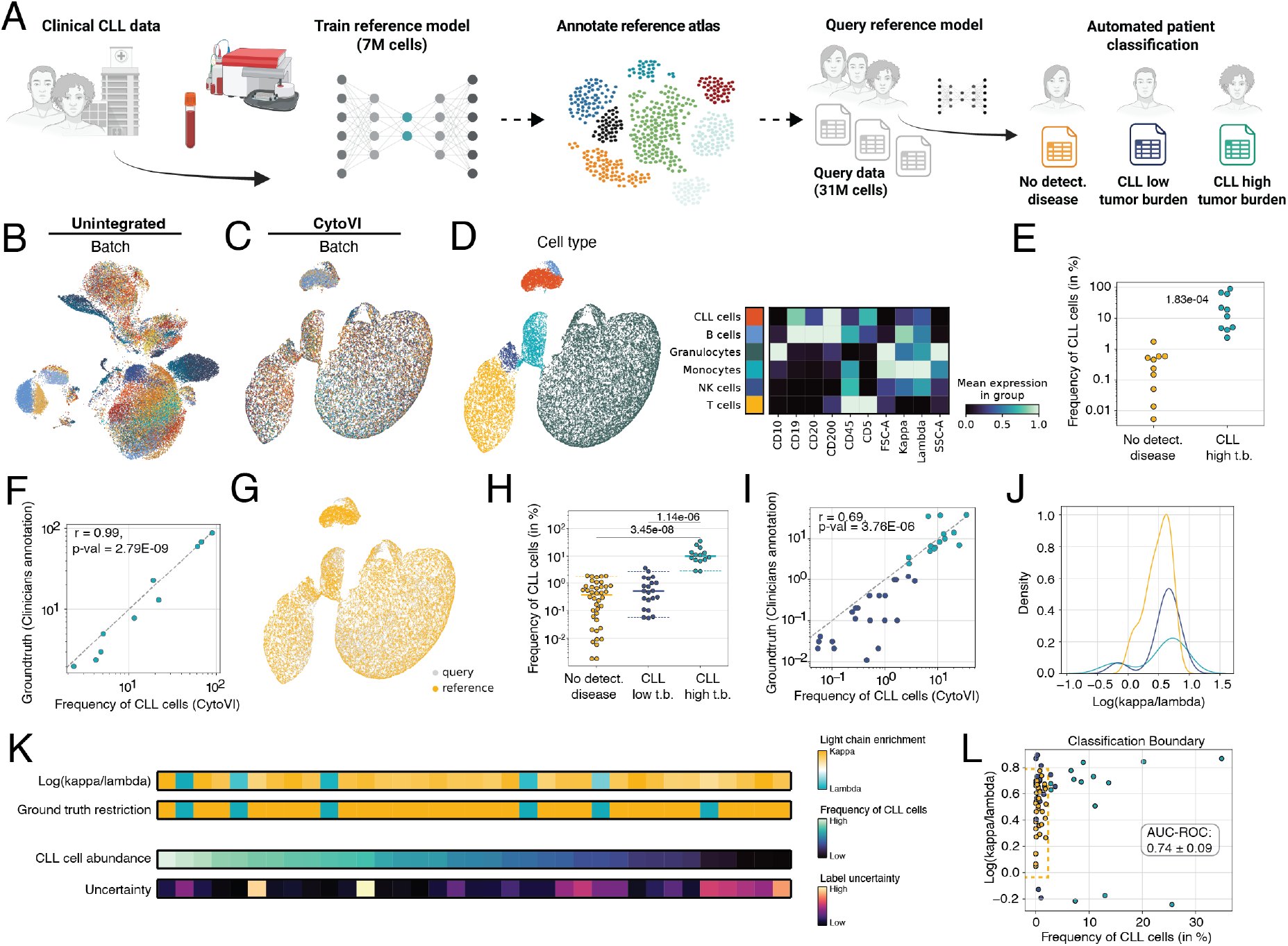
CytoVI guides the automated diagnosis of hematological malignancies. **A:** Schematic overview of the transfer learning task to automatically classify flow cytometry data of chronic lymphocytic leukemia (CLL) patients using CytoVI. **B:** UMAP of the unintegrated data space of flow cytometry samples of PBMCs of ten CLL patients and ten no detectable disease controls used to train the reference model colored by batch (= individual patient). **C:** UMAP of the CytoVI latent space of the same reference data controlling for the observed batch covariate. **D:** UMAP of the CytoVI latent space of ten CLL patients and ten no detectable disease controls (comprising the reference dataset) colored by the corresponding cell type labels obtained from clustering the latent space (left panel). Heatmap showing the protein expression profile of the indicated cell types (right panel). **E:** Scatter plot showing the relative abundance of CLL cells obtained from clustering the CytoVI latent space for CLL high tumor burden patients (teal; N = 10) and patients without detectable ground truth disease in the reference dataset (yellow; N = 10) determined by manual gating of an expert hematologist. The p value corresponds to a Mann-Whitney U test. **F:** Scatter plot for the relative abundance of CLL cells per patient (N = 10) obtained from clustering the CytoVI latent space in the reference dataset compared the ground truth annotation obtained by expert gating. Statistics refer to Pearson’s correlation. **G:** UMAP of the CytoVI latent space displaying the reference dataset (yellow) and additional unseen query data (grey) that was mapped to the latent space using a forward pass of the trained reference model. **H:** Scatter plot showing the relative abundance of CLL cells automatically obtained from mapping new unseen query data into the CytoVI latent space and consecutive label imputation. Frequencies are shown for CLL high tumor burden patients (teal; N = 14; comparable to newly diagnosed CLL patients), CLL low tumor burden patients (blue; N = 21; predominantly minimal residual disease patients) and patients without detectable ground truth disease in the query dataset (yellow; N = 40). Categories on the x-axis correspond to the manual annotations from the hematologist. P values corresponds to pair-wise Mann-Whitney U test in relation to the no detectable disease controls. **I:** Scatter plot for the relative abundance of CLL cells per patient automatically obtained from mapping new unseen query data into the latent space with consecutive label imputation compared to the ground truth annotation obtained by expert gating. Statistics refer to Pearson’s correlation. **J:** Kernel density estimate plot for the Log(kappa/lambda) light chain expression intensity of the automatically detected CLL populations per patient separated by the ground truth annotation into CLL high tumor burden patients (teal), CLL low tumor burden patients (blue) and patients without detectable disease (yellow). **K:** Heatmap displaying the Log(kappa/lambda) light chain expression intensity, ground truth light chain restriction, relative frequency of CLL cells and the uncertainty of the label imputation for the automatically detected CLL cells per patient (N = 35). Each column of the heatmap corresponds to a patient. **L:** Scatter plot displaying the Log(kappa/lambda) light chain expression intensity in relation to the relative frequency of CLL cells for the automatically detected CLL cells per patient (N = 35 CLL patients; N = 40 no-detectable disease controls). The dashed line depicts the decision boundary of a decision tree classifier trained on both features. Statistics refer to mean and standard deviation of the area under the curve (AUC) of the receiver operating characteristic obtained by 5-fold cross validation.

### CytoVI imputes morphological features from mass cytometry data

One inherent limitation of mass cytometry compared to conventional flow cytometry is that morphological features, allowing to draw conclusions about cells’ size or granularity, are absent in mass cytometry. However, these morphological properties can be crucial for biological interpretation, such as the detection of aberrant cells in the context of hematological disorders or provide insights about a cell’s activation state.

To evaluate whether we could utilize CytoVI to impute forward-scattered light (FSC; a proxy for cell size) and sideward-scattered light (SSC; a proxy for cell granularity) from mass cytometry data, we integrated flow and mass cytometry data from healthy PBMCs [14, 15] **(Figure 4A)**. CytoVI controlled for the technical variation between the two distinct technologies and yielded a technology-harmonized latent space that maintained biological variability **(Figure 4B, Figure S6A)**. Employing CytoVI’s generative model, we imputed FSC and SSC in the mass cytometry data. Biaxial visualization of the light scatter features imputed in the mass cytometry dataset yielded typical scatter plots associated with conventional flow cytometry with the relatively large and more granular myeloid fraction indicated by high values for both, FSC and SSC **(Figure 4C, Figure S6B)**. Moreover, we observed that memory CD4 and CD8 T cells displayed consistently higher FSC and SSC values, corresponding to increased cell size and granularity, compared to their naïve counterparts **(Figure S6B)**. These morphological alterations have been described by others before and were associated with biophysical processes underlying T cell differentiation, such as biogenesis and remodeling of mitochondria [39]. Next, we compared the morphological features imputed in mass cytometry data to the ground truth measured in the flow cytometry dataset. This comparison demonstrated accurate imputations of FSC and SSC across the evaluated immune populations **(Figure 4D, Figure S6C)**. In conclusion, we demonstrated that CytoVI can be utilized to integrate antibody-based single cell data across different technologies with consecutive imputation of features that are available in only one of the technologies.

### CytoVI automates the discovery of T cell states associated with nodal B cell lymphomas

The latent space of CytoVI provides a stochastic representation of the state of each cell. This representation facilitates a principled analysis of differences in the composition of cell states between different conditions. Given a set of samples distinguished by a certain covariate (e.g. the presence of disease), this can be achieved by aggregating the posterior distribution of all cells in a sample and then calculating the ratio of aggregated probabilities between different sample groups. The resulting ratios enable a high-resolution differential abundance analysis that does not require *a priori* partition of the cells into groups **(see Methods)**.

To investigate, whether these differential abundance scores could be utilized to automatically detect cells associated with a given disease, we analyzed flow cytometry data of T cell infiltrates in lymph node samples of a large cohort of 63 B cell non-Hodgkin lymphoma patients [40] **(Figure 4E)**. This dataset comprised patients of various nodal B cell lymphomas such as diffuse large B cell lymphoma (DLBCL), marginal zone lymphoma (MZL), mantle cell lymphoma (MCL), follicular lymphoma (FL) as well as resected control lymph nodes (rLN). The dataset was generated using two partially overlapping flow cytometry antibody panels (12 proteins each, with an overlap of 8 proteins), which were analyzed separately by the study authors using manual gating to identify T cell states associated with the respective disease entity. Using CytoVI we integrated the flow cytometry data of the two panels and imputed non-overlapping proteins **(Figure 4F, Figure S7A and B)**.

The publicly available flow cytometry dataset from *Roider et al*. did not comprise cell state annotations from the original study. To enable a cross-comparison of T cell states detected using CytoVI’s label-free differential abundance functionality with the T cell clusters defined by the original study authors (using manual gating), we clustered the latent space and used the original author’s annotation strategy to divide cells into similar T cell subsets [40] **(Figure S7C)**. Next, we computed CytoVI’s label-free differential abundance scores for each T cell to identify cell subsets that are enriched in the respective disease entities. This analysis highlighted regions in latent space that are specifically enriched (red) or depleted (blue) for the given disease entity **(Figure 4G)**. Concatenating these scores with the coordinates of each cell in latent space, followed by graph-based clustering, revealed T cell states with differential abundance across the interrogated lymphoma entities **(Figure 4H, Figure S7D)**. Notably, the frequency of these differential abundance T cell states was highly similar for cells measured across the two distinct antibody-panels, validating the applicability for an integrated analysis of both panels simultaneously (ICC = 0.99, p = 5.97e-12; **Figure S7E**).

Next, we matched the annotations based on the confusion matrix with the T cell states from *Roider et al*.[40] **(Figure S7F)**. In line with the original authors observation, we found a cluster of naïve CD8 T cells and a cluster of naïve CD4 T cells to be enriched in the control lymph nodes (rLN) **(Figure 4I, Figure S7F and G)**. Lymph node samples originating from FL patients demonstrated increased abundance of a cluster of follicular T helper cells, while samples from DLBCL and MZL patients were enriched for a cluster of proliferating T cells **(Figure 4I, Figure S7G)**. Yet, opposed to the report from the original authors describing a general proliferating T cell subset, we found that the proliferating T cell cluster enriched in DLBCL and MZL patients comprised predominantly proliferating CD8 T cells and to a lesser extent proliferating T helper cells, highlighting the potential of the integrated label-free differential abundance analysis to detect rare cell states associated with disease **(Figure 4G-I, Figure S7B and G)**. Interestingly, our label-free differential abundance analysis detected a small cluster of CD4/CD8 double-negative T cells that was specifically enriched in DLBCL and MCL patients and was not reported by *Roider et al*. [40] **(Figure 4I)**. These double-negative T cells displayed characteristics of activated cytotoxic memory T cells (KLRG1^+^, CD31^+^, CD69^+^, PD1^+^, and TIM3^+^) but the protein markers included in the two antibody panels did not allow a more confident assignment of these cells to a specific T cell subset **(Figure S7B)**.

Thus, we proceeded with integrating the flow cytometry data of the BNHL cohort with CITE-seq data of a subset of the patients, impute the transcriptome for T cells in the flow cytometry dataset and perform a differential gene expression analysis of the double-negative T cell cluster to investigate the lineage identity of this cluster **(Figure 4J)**. Computing the confusion matrix between cell type labels of the flow cytometry dataset and the nearest neighbor in the CITE-seq dataset based on the integrated CytoVI latent space indicated good cell type preservation of the integrated latent space for most cell types between the two technologies **(Figure S7H)**. To gauge the extent to which a cell’s transcriptome could be imputed from paired CITE-seq data, we computed the correlation between protein expression measured by flow cytometry and the imputed gene expression of the corresponding transcripts. We observed strong concordance between measured protein levels and imputed transcript expression, including for proteins not present in the CITE-seq panel, such as the proliferation marker Ki67 and the transcription factor FOXP3 **(Figure 4K, Figure S7I and J)**. Of note, CD69 demonstrated a negative correlation between measured protein expression and imputed gene expression of its corresponding transcript **(Figure S7J)**. Such negative correlation was also observed in the CITE-seq dataset alone and documented by others in two independent CITE-seq datasets indicative of a poor relationship between protein and transcript expression of CD69 **(Figure S7K)** [41]. Overall, comparison of the protein-gene expression relationships observed in the CITE-seq data in relation to the imputed RNA-protein relationship in the flow cytometry data confirmed that imputation effectively preserved these associations **(Figure S7J and K)**.

To determine the lineage identity of the infiltrating double-negative T cell subset in DLBCL and MCL patients, we performed differential gene expression analysis of the imputed transcriptome data (comparing the double-negative subset to all other T cells in our flow cytometry data). Consistent with our previous observations, the double-negative T cell subset exhibited elevated expression of cytotoxic effector molecules (*GZMA, GZMB, PRF1, GZMK*) and expressed transcripts for both, activating (*NKG7*) and inhibitory (*KLRC2, KLRC3, KLRG1*) innate-like receptors associated with cytotoxicity **(Figure 4L and M, Figure S7B)**. Notably, the T cell subset lacked expression of the T cell receptor (TCR) *β* chain but expressed transcripts encoding for the constant chains of TCR *γ* and *δ* constant chains, strongly indicative of an affiliation to the *γδ* T cell lineage **(Figure 4L and M, Figure S7L)**. Moreover, the *γδ* T cell subset demonstrated weak enrichment of *TRDV1*, encoding the V*δ*1 TCR chain, and expressed the cytokine *IFNG* **(Figure 4L and M, Figure S7M)**. These IFN-*γ*-producing cytolytic V*δ*1 memory T cells have been previously observed in tumors, where they mediate anti-tumor immunity through innate-like receptors, independent of cognate peptide-MHC recognition [42, 43]. Interestingly, these V*δ*1 memory T cells have been reported to be enriched in the blood and tumors of patients with DLBCL and were suggested as immunomodulatory targets for therapy [44]. To validate the gene signature of the *γδ* T cell subset in the B cell non-Hodgkin lymphoma patient cohort, we next analyzed the measured transcriptome of T cells in BNHL patients detected by CITE-seq. In line with the imputed transcriptome of the double-negative T cell cluster detected by flow cytometry, we identified a *γδ* T cell cluster in the CITE-seq data that was characterized by the production of *IFNG* and cytotoxic molecules and expressed innate-like receptors associated with cytotoxicity **(Figure 4N, Figure S7M)**. Similarly as observed in the flow cytometry data, there was a trend for an enrichment of the *γδ* T cell subset in lymph nodes of DLBCL and MCL patients albeit not statistically significant due to the low number of cells and patients that could be analyzed using CITE-seq **(Figure S7N)**. Collectively, we demonstrated CytoVI’s ability to automatically identify disease-associated cell states. By integrating high-throughput technologies like flow cytometry with transcriptome-wide analyses using CITE-seq, CytoVI combined the advantages of these complementary technologies, the throughput of cytometry with the dimensionality of single cell genomics, and enabled a powerful exploratory characterization of cellular states for preclinical studies.

### CytoVI facilitates the automated diagnosis of chronic lymphocytic leukemia patients

Beyond the role as an exploratory single-cell technology for preclinical research, flow cytometry is widely used in clinical practice to guide treatment decisions and diagnose immune-related disorders, such as hematological malignancies and primary immunodeficiencies. However, in a clinical setting, data are typically generated on a per-patient basis and must be analyzed individually using manual gating due to technical variation between measurements. Given CytoVI’s ability to distinguish biological from technical variation, we next evaluated its potential to facilitate joint analysis and automate the diagnosis of hematological malignancies using transfer learning **(Figure 5A)**. To this end, we collected clinical flow cytometry data from chronic lymphocytic leukemia (CLL) patients using standardized antibody panels designed for clinical monitoring of tumor burden at the diagnostic hematology unit of the University Hospital in Basel (**Supplementary Table S1**). According to the diagnostic procedures, PBMCs from each patient were analyzed by an expert hematologist and classified into three distinct categories based on the abundance of CLL cells: 1) no detectable disease, where no CLL cells were identifiable in the blood; 2) high tumor burden CLL, defined as cases, where CLL cells were present in *>* 2% of total PBMCs, corresponding to the expected clinical phenotype of newly diagnosed patients; and 3) low tumor burden CLL, where CLL cells were present *<* 2% of the total PBMCs, reflecting minimal residual disease following therapy.

To construct a reference model for automated analysis of frequencies of cell subsets followed by diagnosis of incoming (unseen) patient samples, we selected a dataset comprising ten patients with no detectable disease and ten patients with high tumor burden CLL. Despite the use of standardized clinical flow cytometry panels, batch effects arising from technical variation between experiments hampered a joint downstream analysis **(Figure 5B, Figure S8A)**. However, by explicitly accounting for technical variability, CytoVI generated an integrated latent space that mitigated technical variation and preserved only biological variation **(Figure 5C, Figure S8B)**. Graph-based clustering of the latent space identified the major leukocyte subsets in the peripheral blood, including CLL cells, characterized by a CD5^+^ CD20^dim^ phenotype **(Figure 5D)**. Quantification of CLL cell frequencies per patient enabled a clear distinction between patients with no detectable disease and those with high tumor burden, with relative CLL cell abundance strongly correlating with clinical annotations **(Figure 5E, F, Figure S8C)**. This confirmed that CytoVI could be employed to mitigate technical variation in clinical flow cytometry data and facilitate a combined analysis. We then leveraged the trained model, along with its corresponding cell type annotations, as a reference for analyzing additional flow cytometry samples that were obtained at the diagnostic hematology unit, but not seen during training (query samples). Encoding these query flow cytometry data into the reference model’s latent space (leveraging the scArches implementation in scvi-tools [45, 10]) demonstrated that incoming samples could be effectively mapped, while mitigating technical variation through a simple forward pass **(Figure 5G)**. Next, we automatically assigned cell type labels for the query cells based on a majority vote among their nearest labeled neighbors in latent space **(see Methods)**. We observed that for most query cells, the majority of neighboring reference cells have been assigned with the same label (hence, no added uncertainty due to KNN), with the exception of a small fraction of query cells that were projected near the interface between CLL cells and healthy B cells **(Figure S8D)**. Nevertheless, quantification of the relative abundance of imputed CLL cells in the query samples allowed for clear distinction between samples taken from high tumor burden CLL patients (as determined manually) and control samples with no detectable disease **(Figure 5H, S8E)**. While the automatically detected CLL cell frequency alone was insufficient to distinguish low tumor burden CLL patients (who may harbor very small residual CLL clones during therapy) from no detectable disease controls, the automatically detected frequency of CLL cells correlated well with manual hematologist annotations and was sufficient to distinguish between high and low tumor burden **(Figure 5H-I, S8E)**. Thereby, we demonstrate that CytoVI can be utilized in a transfer learning setting to automate the diagnosis of unseen CLL patients once they are presented to the clinic.

A hallmark of CLL cells is their monoclonality, indirectly characterized by restriction to either *κ* or *λ* light chain expression, an important clinical feature, which can be used to track clonal dynamics over time. To include this parameter in our automated report, we estimated the *κ*-to-*λ* light chain ratio of the uncorrected protein expression data in the automatically classified CLL cells. As expected, CLL patients with detectable clones exhibited skewed *κ*-to-*λ* ratios indicative of a monoclonal population, whereas patients classified as having no detectable disease displayed a more balanced distribution **(Figure 5J)**. Comparison of automatically calculated *κ*/*λ* light chain restriction with annotations assigned manually by a hematologist demonstrated strong concordance **(Figure 5K)**. For one single case the light chain restriction was misclassified by the automatic procedure. In this patient the CLL population was exceptionally small, and the model indicated high uncertainty for the label imputation **(Figure 5K)**. Finally, integrating both the predicted relative frequency of CLL cells and the *κ*-to-*λ* ratio into a decision tree classifier improved classification performance, enabling robust differentiation of CLL patients — including those with minimal residual disease (AUC-ROC: 0.74 ± 0.09), which remains challenging to detect from no detectable disease controls even for expert hematologists **(Figure 5K-L, S8F)**. These findings highlight the potential of CytoVI to automate and standardize flow cytometry based diagnostics via transfer learning, while controlling for technical variability and providing uncertainty estimates about predictions. This approach could support clinical decision-making by providing a scalable, objective, and reproducible framework for monitoring disease progression and treatment response in hematological malignancies.

## Discussion

The increasing number of cytometry datasets available in the public domain presents great opportunity to enhance our understanding of cellular systems and gain insight about diseases at an unprecedented scale. However, these datasets are currently analyzed in isolation due to the lack of computational methods capable of standardizing and integrating information across technologies, studies, and experimental conditions. This limitation hampers our ability to accumulate knowledge by jointly analyzing data from distinct studies or when distinct antibody panels or technologies are employed. Moreover, when longitudinal sampling is necessary but sample cryo-preservation may be impracticable, technical variation often hinders a joint analysis. To address these challenges, we introduce CytoVI, a probabilistic generative model designed for the integrative analysis of antibody-based cytometry data. CytoVI is incorporated into scvi-tools [10] and connects well with Scanpy, a popular single cell analysis framework from single cell genomics [46]. Thereby, CytoVI provides a highly scalable and effective end-to-end pipeline for the computational analysis of cytometry data. We demonstrated that CytoVI can effectively handle a broad array of integration scenarios, including those comprising distinct cohorts, antibody panels, and cytometry technologies. By embedding cells into an informative latent space, imputing missing protein measurements, predicting gene expression (via integration with CITE-seq), identifying differentially expressed proteins or differentially abundant cell states, and conducting automated annotations, CytoVI provides a statistically rigorous framework for harmonizing diverse cytometry datasets. These capabilities enable researchers to extract meaningful biological insights that would otherwise be inaccessible when working with isolated datasets. Apart from its utility in exploratory preclinical studies, we have demonstrated the applicability of CytoVI in clinical settings, where flow cytometry is prevalent for diagnostic procedures and patient monitoring. We showcased that CytoVI can be applied to automate the diagnosis of CLL patients using clinical flow cytometry screens, which can be challenging as patients are measured individually and the resulting flow cytometry data contains a mixture of technical and biological variation. Thereby, CytoVI addresses one of the most pressing challenges of clinical flow cytometry as outlined by the EuroFlow Consortium for clinical cytometry [47]. CytoVI was capable of reliably identifying CLL cells along their associated routine clinical parameters (such as immunoglobulin light chain restriction) in the blood of patients up to a detection limit that is relevant for diagnostic procedures, thereby standardizing the clinical analysis and sparing personal and financial resources at clinical centers. One key advantage of our probabilistic approach is its intrinsic ability to estimate uncertainty, which enhances interpretability and confidence in critical clinical decision-making. To ensure accessibility for clinical centers that are not equipped with the specialized hardware infrastructure, CytoVI could be deployed to clinical centers via cloud-based solutions. Alternatively and to mitigate patient privacy concerns, pretrained models could be distributed and accessed locally using regular desktop computers for fast and efficient inference in transfer learning settings.

In preclinical studies, high-dimensional cytometry is often used as an exploratory hypothesis-generating technology to interrogate large cohorts of precious patient samples with a predefined panel of protein markers. Due to the destructive nature of the technology, the retrospective analysis of additional proteins is experimentally impossible. We have demonstrated that CytoVI can be utilized in these cases to retrieve probabilistic expression estimates of proteins not measured in the original study. This functionality extends the utility of archived datasets, allowing researchers to generate new insights from existing data. Moreover, we have demonstrated that CytoVI can be utilized to combine the advantages of high-throughput technologies such as flow cytometry with the benefits of single cell genomics. The integrative analysis of flow cytometry and paired CITE-seq data facilitated an unbiased characterization of cellular states across the entire transcriptome, while maintaining the possibility to analyze millions of cells. We showcased that the combination of these highly complementary technologies could uncover a rare *γδ*T cell subset in tumors of DLBCL patients that would have been undetectable using either of these technologies alone. As novel antibody-based single-cell technologies emerge, offering interrogation of novel modalities to measure cellular states, we anticipate that CytoVI will be capable of integrating these new technologies. For instance, Molecular pixelation enables subcellular quantification for protein cooccurrences within a cell in addition to protein abundance estimates, which could be seamlessly incorporated into CytoVI’s framework, further expanding its applicability [48]. InfinityFlow, the combination of machine-learning with plate-based screens containing lyophilized antibodies targeting more than 300 proteins via flow cytometry has demonstrated marked potential for the generation of large-scale protein atlases using traditional cytometry [7]. CytoVI could empower these largescale plate-based protein screens and enable community-accessible organ-scale protein atlases that can be queried using transfer learning approaches. By leveraging these reference datasets, CytoVI can generate hundreds of stochastic protein expression estimates per cell, significantly enriching downstream analyses of newly generated data. The ability to query large-scale cytometry data through probabilistic models represents a transformative step toward standardizing cytometry data interpretation and application. We envision CytoVI as a first step towards a model-centric view for cytometry data, analogous to recent advances in other domains. By continually learning from expanding cytometry datasets, we anticipate that these generative models improve on generalization, robustness, and predictive power across studies and clinical applications. These models could support the development of intelligent, data-driven systems for immune profiling, biomarker discovery, and personalized medicine, fundamentally transforming the way cytometry data is utilized in both, preclinical research and clinical settings.

## Methods

### The CytoVI model

CytoVI is a probabilistic latent variable model for antibody-based single cell technologies, such as flow cytometry, mass cytometry or CITE-seq. Data from each of these technologies can be represented as an expression matrix *X* ∈ ℝ^*N ×P*^, in which each element *x*_*np*_ denotes the expression of protein *p* for cell *n* among *P* total proteins and *N* total cells. In flow and mass cytometry protein expression data is generated via indirect labeling of antibodies that are conjugated to fluorophores or isotopically pure heavy metal ions. Thus, detected protein intensities are relative measurements that heavily depend on the choice of respective reporter molecules coupled to the respective antibody. To account for the relative nature of antibody-based single-cell data and enhance its suitability for visualization and downstream analysis, cytometry data is typically preprocessed using hyperbolic arcsin, logicle, or biexponential transformations and feature-wise scaling to yield comparable marker expression ranges [12, 13]. To provide flexibility for various applications and common downstream tasks, CytoVI accommodates these preprocessing steps, allowing for the incorporation of any of these transformations into the model. Specifically we assume that *x*_*np*_ is a transformed and/or scaled protein expression matrix. Additionally, each cell can be associated with additional observed sample-level covariates that can be controlled for in CytoVI’s latent representation, such as categorical nuisance factors, representing experimental batches, utilized technologies or antibody panels applied. For simplicity we will describe the case of a one-hot encoded batch identifier *s*_*n*_ ∈ {1, …, *S*}. We assume that the protein expression *x*_*np*_ for cell *n* and protein *p* can be generated from a *d*-dimensional latent vector *z*_*n*_, reflecting a cell’s intrinsic state. CytoVI utilizes a variational autoencoder (VAE) to approximate the posterior distribution *p*(*z*_*n*_|*x*_*n*_).

### The generative model

We begin by describing the generative model of CytoVI. We assume an isotropic Gaussian prior or a mixture of Gaussians (1) with a diagonal covariance matrix for the latent space *z*, where 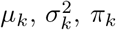 are learnable parameters and denote the mean, the standard deviation and the mixing coefficient for the *k*-th component, respectively:

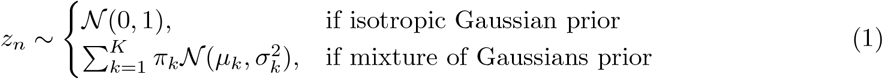

The choice of the prior depends on the expected biological variation across cells in the corresponding dataset. Homogeneous datasets are best modeled using the isotropic Gaussian prior, whereas the mixture of Gaussian prior is best applied in heterogeneous samples comprising various different cell types. By default CytoVI models the prior for the latent space *z*_*n*_ as a mixture of Gaussians, which increases the expressiveness of the latent space and has excelled for the integration of large single cell studies [11]. When prior knowledge about the cells present in the dataset is available (e.g. cells in the dataset have been annotated before), the prior distribution can be conditioned on these labels to weakly inform the latent representations. Specifically, we adjust the mixture component weights by incorporating a one-hot encoded label vector *y*, such that:

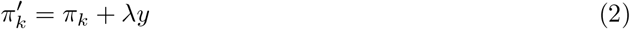

In this case, *λ* is a hyperparameter that determines the weight for the existing cell type label on the prior (default *λ* = 10).

The generative network models the observed protein expression *x*_*np*_ as a function of each cell’s latent vector *z*_*n*_ and its corresponding one-hot batch vector *s*_*n*_:

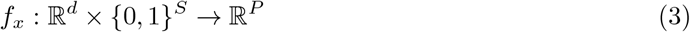

Here, *f*_*x*_ denote the non-linear mappings generated by the decoder network. Finally we describe our observation model using either a Gaussian, parameterized by the mean *µ*_*np*_ and standard deviation 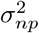, or beta distribution, which is parameterized using the shape parameters *α*_*np*_ and *β*_*np*_ (4):

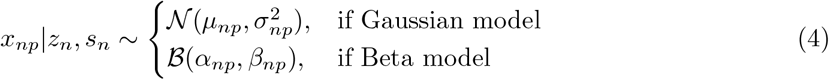

The choice of the protein likelihood distribution is determined by the specific preprocessing steps applied to the data. By default, we employ a Gaussian emission distribution to model cytometry data that has undergone arcsinh or log-transformation. In cases where the data has been scaled to a [0, 1] range with or without prior transformation, a beta distribution can be utilized to model the bound values.

#### Variational approximation and training procedure

Since the posterior distribution *p*(*z*_*n*_|*x*_*n*_) is intractable, we rely on variational inference to learn a posterior approximation *q*(*z*_*n*_ | *x*_*n*_), similar to our previous work [49]. The variational posterior is chosen to be a Gaussian with a diagonal covariance matrix, with parameters given by a Multi-Layer Perceptron (MLP) encoder *f*_*z*_ applied to *x*_*n*_.

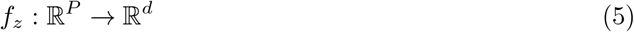

During training the evidence lower bound (ELBO) is optimized with respect to the inference and generative network parameters using mini-batch stochastic gradient descent. The ELBO consists of a reconstruction term, which encourages the reconstruction of the observed data *x* given *z*, and the Kullback-Leibler (KL) divergence term that regularizes the posterior distribution:

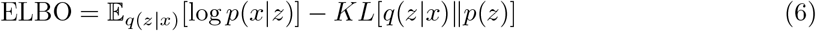

#### Handling of overlapping antibody panels

Integration of antibody-based single cell experiments is often constraints by different input features, e.g. integration of two flow cytometry datasets employing different antibody panels, resulting in missing values in the expression matrix *X* after concatenation. We therefore apply a masking scheme for the unobserved proteins similarly to what was proposed in our previous work [17].

Let 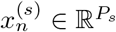 represent the observed input features for cell *n* in batch *s*, where *s* = 1, …, *S*, and *P*_*s*_ denotes the number of features in batch *s*. The panel set of each batch is denoted 𝒯 _*s*_, such that | 𝒯 _*s*_| = *P*_*s*_. We define the set of shared features among all batches as ℐ, where:

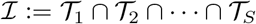

In these cases the encoder network processes only the shared features ℐ to compute *z*_*n*_:

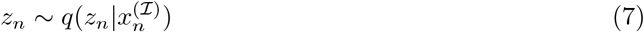

The decoder network reconstructs the union of the features for each of the batches denoted as 𝒰:= 𝒯 _1_ ∪ 𝒯 _2_ ∪ · · · ∪ 𝒯 _*S*_. This yields a reconstructed protein expression vector 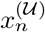 for each cell. In order to handle missing values during training, we generate binary feature masks, 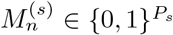 for each batch *s* = 1, …, *S*, where:

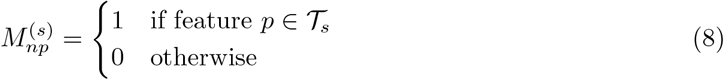

We utilize 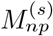 to mask the reconstruction loss for features *p* that were not observed in the respective batch.

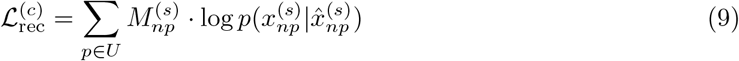

To encourage latent space mixing in case of overlapping antibody panels, we include an adversarial classifier that penalizes the discrimination of cells between employed antibody panels, as proposed in TotalVI [17].

#### Imputation of missing proteins

When integrating data generated using distinct antibody panels, CytoVI can impute missing proteins. Specifically, if a protein *p* was not measured in batch *s* but was observed in batch *s*^*′*^, we perform counterfactual decoding to infer its expression.

To achieve this, we first sample the latent representation *z*_*n*_ from the approximate posterior:

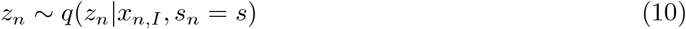

where *x*_*n,I*_ represents the subset of features shared across batches. Using this latent representation, we generate the missing protein’s expression by decoding while conditioning on batch *s*^*′*^:

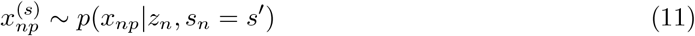

This procedure reconstructs the missing protein expression in batch *s* using knowledge from batch *s*^*′*^. A similar counterfactual decoding can also be applied to data that has been generated using the same antibody panel in order to obtain a stochastic batch-corrected protein expression estimate, analogous to our previous work for single cell transcriptomics [49].

#### Implementation of CytoVI

CytoVI is implemented as a PyTorch-based model within the scvi-tools framework [10]. While the model exposes hyperparameters for customization, it utilizes a default encoder-decoder architecture, each consisting of a single hidden layer with 128 hidden units. Model parameters are optimized using the Adam optimizer with weight decay. The dimensionality of the latent space *z*_*n*_ is primarily influenced by the biological characteristics of the sample, as well as the number *P* and quality of protein markers in the dataset. Based on our observations, latent space dimensions *d* in the range of 10 to 20 typically yield optimal results for batch correction while preserving biological variation. The default heuristic for determining the latent space dimension is and constrained between 10 and 20. Additionally, we set the number of components *K* in the mixture of Gaussian prior to match the dimension of latent space *d* by default.

### Differential protein expression analysis

CytoVI leverages the trained generative model for testing differentially expressed proteins using the Bayesian framework proposed earlier [50]. For this we consider cells to be stratified into categories *c*_*n*_ that represent our covariate-of-interest (e.g. disease state, response to therapy). To identify differentially expressed proteins between cells of the group *c*_*a*_ versus *c*_*b*_, we first perform uniform sampling with replacement of the posterior distribution of cells among each group. We then estimate the log fold change (LFC) 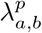 in generated expression 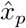 for protein *p* between pairs of cells *a* and *b* belonging to groups *c*_*a*_ and *c*_*b*_:

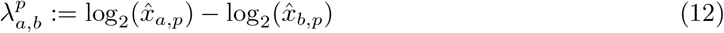

We then estimate the probability that protein *p* is differentially expressed between cells of *c*_*a*_ and *c*_*b*_ by assessing the fractions of posterior samples for which *λ*^*p*^ is larger than a predefined effect size threshold *δ* (defaults to 0.25).

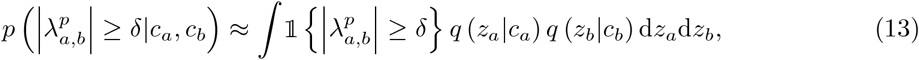

To account for batch effects in differential expression (DE) estimation, we perform counterfactual decoding across all possible batch conditions *s*_*n*_ and compute the DE probability as the average over these counterfactual samples.

Since cytometry data is often feature-wise normalized to the [0, 1] range, we apply numerical clipping to expression estimates at the boundaries. Specifically, values of exactly 0 or 1 are adjusted to 0 + *ϵ* and 1 − *ϵ*, respectively, where *ϵ* = 10^−6^, to ensure numerical stability in downstream computations.

### Label-free differential abundance analysis

CytoVI incorporates a label-free differential abundance estimate across conditions of interest that can be utilized to identify cellular states associate with a sample covariate of interest (such as presence of disease) as described in [11]. For this we aggregate the approximate posterior distribution *q*_*v*_(*z*|*x*) for each sample *v* across *n*_*v*_ cells:

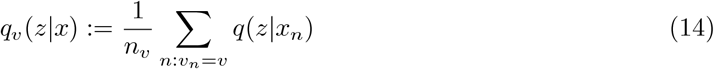

Accordingly, we aggregate the posterior densities across all samples of a given covariate in *A* ⊂ {1, …, *V*_*a*_} and *B* ⊂ {1, …, *V*_*b*_} into *q*_*A*_(*z*) and *q*_*B*_(*z*) and estimate the ratio of enriched and depleted regions in latent space with respect to the categorical covariate:

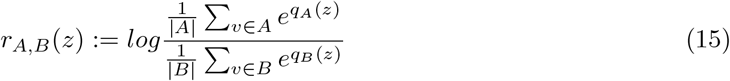

The resulting differential abundance scores *r*_*A,B*_(*z*) can be utilized for a cluster-free analysis of compositional alterations associated with the covariate of interest.

#### Label transfer between query and reference datasets

CytoVI facilitates transfer learning by leveraging pre-trained reference models, following the base implementation of the scArches class in scvi-tools [45, 10]. Unseen query data can be projected into the latent space of a trained reference model, enabling batch-corrected embeddings without additional training. To transfer annotations, we perform label propagation via nearest neighbor search in the shared latent space. Specifically, for each query cell, we identify its k-nearest neighbors (default k = 20) in the reference dataset using the approximate nearest neighbor search algorithm implemented in PyNNDescent. The query cell is then assigned the most frequent label among its neighbors, with the relative abundance of the majority label serving as an uncertainty estimate.

#### Imputation of missing modalities between query and reference datasets

CytoVI’s latent representation enables the imputation of additional modalities that were not observed during training. We demonstrate its applicability by imputing transcriptomic profiles in flow cytometry data using paired CITE-seq datasets. Analogous to label imputation, we leverage a shared latent representation of protein expression across both modalities by training a CytoVI model that accounts for technical variability between the technologies. To impute continuous covariates, such as normalized RNA expression from CITE-seq data, we identify for each cell in the flow cytometry data its k-nearest neighbors (default k = 20) in the protein latent space of the CITE-seq dataset using pyNNDescent. The imputed gene expression vector for the query cell is then obtained by averaging the expression profiles of its 20 nearest neighbors in the reference dataset.

### Model evaluation using flow cytometry, mass cytometry and CITE-seq data

#### Analysis of fluorescence intensities in homogeneous microspheres

To determine a suitable distribution for the protein likelihood we analyzed the fluorescent intensities of microsphere that either did not bind fluorescent antibodies at all (negative fraction) or bound a homogenous number of fluorescent antibodies. For this, compensation beads (BioLegend) were used to generate single-stain controls using the following fluorescent antibodies: anti-human CD3 Pacific Blue (Clone UCHT1; BioLegend), anti-human TIGIT BUV395 (Clone 741182; BD Biosciences), anti-human CD56 BV421 (Clone 5.1H11; BioLegend), anti-human CD25 RB780 (Clone 2A3; BD Biosciences). For each stain, one drop of compensation beads was mixed with one drop of negative bead controls, followed by the addition of 200 *µ*L of PBS. From this mastermix, 50 *µ*L were incubated with 0.5 *µ*L of the respective antibody for 15 minutes at 4°C. After incubation, 200 *µ*L of PBS was added, and the samples were centrifuged at 500 × g for 5 minutes. The supernatant was discarded, and the samples were resuspended in 200 *µ*L of PBS before acquisition on a Cytek Aurora 5L spectral flow cytometer using default settings. Beads were identified based on FSC and SSC using biaxial gating in FlowJo and positive and negative fraction of antibody-bound beads were exported separately for each staining condition. The resulting.fcs files were imported into Python using the readfcs library. Quantile plots were drawn using Scipy.stats.

#### Flow cytometry data of healthy donor PBMCs

To evaluate the distribution of fluorescence intensities in flow cytometry data, we analyzed full spectrum cytometry data of cryopreserved PBMCs of a healthy donor that served as the normalization control in a recent SARS-CoV-2 vaccine study [14]. The flow cytometry dataset was down-sampled to 100 000 cells and comprised 35 protein parameters and additional morphological features for FSC and SSC. Doublets, dead cells and fluorescent aggregates have been removed using manual gating in FlowJo. The exported.fcs files have been imported into Python using the readfcs library and stored as an AnnData object, similarly as employed for single cell genomics data. The raw fluorescent intensities of the protein markers (excluding scatter features) were transformed using a hyperbolic arcsin transformation with a global co-factor of 2000 using CytoVI’s preprocessing module. The hyperbolic arcsin transformed expression values (protein markers and scatter features) were scaled feature-wise to a range of [0, 1]. We next computed the nearest neighbor graph based on the input feature representation using Scanpy [46]. The resulting neighbor graph was used to compute UMAP and subsequently subjected to Leiden clustering using Scanpy. Leiden clusters have been manually annotated into the most abundant leukocyte populations in the peripheral blood based on key protein marker expression profiles. For the benchmarking of batch integration performance, we utilized a technical replicate of the same healthy donor that was cryopreserved and analyzed by a full spectrum analyzer in a different batch, serving as an internal batch control in the original study [14]. The technical replicate has been processed using the same procedure as outlined above in order to obtain independent annotation of the two replicates into the same leukocyte subsets.

#### Mass cytometry data of healthy donor PBMCs

To benchmark CytoVI’s ability to model mass cytometry data, we accessed mass cytometry data comprised 36 surface and intracellular protein markers of two healthy donor PBMCs that were used as normalization controls in a recent study of monozygotic twin pairs discordant for multiple sclerosis [15]. The two healthy donors were measured in one individual batch using a live cell barcoding approach to mitigate technical variation between the samples. Therefore, the concatenated mass cytometry data of both of the samples was processed in a similar approach as outlined above. In short, the data was down-sampled to 100 000 cells, transformed using a hyperbolic arcsin transformation with a global cofactor of 10 and features-wise scaled to a range of [0, 1]. CD138 was excluded from the analysis due to unspecific staining as carried out by the original study. The nearest neighbor graph has been constructed based on all remaining input features of the mass cytometry data of both concatenated samples and was used for UMAP and consecutive Leiden clustering. PBMCs have been manually annotated into the major leukocyte populations.

#### CITE-seq data of healthy donor PBMCs

To evaluate CytoVI’s ability to model the protein expression measured using CITE-seq, we accessed CITE-seq data across a panel of 228 surface protein markers of PBMCs from an HIV vaccine trial [16]. To evaluate data generated from a single batch and enable a cross-comparison to healthy PBMCs measured by flow and mass cytometry, we selected one individual donor of the CITE-seq dataset at baseline (before any vaccination applied). As the original data contained cell annotations, we utilized these labels as the ground truth cell type annotations. Similarly as for the flow and mass cytometry data, the protein data of the CITE-seq dataset was transformed using a hyperbolic arcsin transformation with a global co-factor of 5 and each feature was scaled to a range of [0, 1].

#### Posterior predictive checks

To assess the extent to which CytoVI is able to generate protein expression data that closely matches the observed data we performed posterior predictive checks. Posterior predictive checks were performed using healthy PBMCs of flow cytometry data from [14], mass cytometry data from [15] and CITE-seq data from [16] as outlined above. Histograms of selected markers measured across flow cytometry, mass cytometry and CITE-seq were drawn using CytoVI’s plotting module. For each technology three separate CytoVI models were trained using either 1) a Gaussian protein likelihood operating on the arcsinh transformed data without additional feature scaling, 2) a Gaussian protein likelihood operating on the arcsinh transformed and feature-wise scaled data and 3) a Beta distribution for the protein likelihood operating on the arcsinh transformed and feature-wise scaled data. For the mass cytometry data a latent space of size *d* = 10 was chosen across each of the models, while we chose a latent space of size *d* = 20 for flow cytometry and CITE-seq data. Posterior predictive checks were performed using an extension of the scvi.criticism module. As a naive baseline, we employed Factor Analysis (FA) using the scikit-learn package, following the implementation described in [17]. For each model, we generated ten posterior predictive samples from 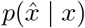 and compared the mean coefficient of variation (CV) across both the cell and protein axes to the CV of the observed data. To quantitatively assess model performance, we computed the mean absolute error (MAE), as well as Pearson’s and Spearman’s correlation coefficients between each model’s generated CV and the observed CV across both, the cell and protein axes.

#### Latent space interpretability

To assess to what extent the CytoVI latent space captured biological variation we utilized the CytoVI models for each of the technologies utilized in the posterior predictive checks. The latent space interpretability was demonstrated for models using a Gaussian protein likelihood that performed best during posterior predictive checks. To analyze the structure of the d-dimensional latent space (*d* = 10 for mass cytometry and *d* = 20 for flow cytometry and CITE-seq), we constructed a k-nearest neighbors graph based on the latent embeddings and applied UMAP using Scanpy. The ground truth cell annotations that were either obtained from the original study if available (for the CITE-seq data) or generated from clustering the data space (see above), were visualized on the UMAP of the latent representation. To assess whether the CytoVI latent space facilitates *de novo* cell typing, we applied Leiden clustering on the latent representation for the flow cytometry data using Scanpy. The ground truth cell annotations and Leiden clusters obtained from the CytoVI latent space were compared by computing the confusion matrix between the two labels using Scanpy and reordering the matrix using a linear sum assignment implemented in Scipy.

#### Benchmarking of integration performance

Benchmarking of integration performance was conducted using the annotated flow cytometry data from cryopreserved PBMC replicates [14]. Cy-toVI was benchmarked against cyCombine and Harmony, two widely used methods for cytometry data integration that do not inherently require normalization controls and are therefore not restricted to experiments, where such controls are available [23, 24]. Integration using cyCombine was performed in R using default parameters. cyCombine’s workflow necessitates an intermediate self-organizing map (SOM) clustering step, followed by batch correction applied to the original arcsinh-transformed protein expression values. To ensure comparability across methods, we postprocessed the cyCombine-corrected expression matrix by applying the same transformation used during label generation — either min-max, z-score, or rank scaling—before downstream analysis. Accordingly, CytoVI and Harmony were also evaluated using each of the pre-processing options (min-max, z-score or rank scaling). Despite CytoVI’s assumption of a Gaussian likelihood to model protein expression data (which is incompatible with rank-scaled data), we included the integration of CytoVI on rank-scaled data for completeness of the comparison. As before, we trained the CytoVI models by optimizing latent space of size *d* = 20 with default parameters. Harmony integration was performed using the Python implementation using default parameters. We evaluated batch correction and biological conservation across integration methods using scvi.scib-metrics, employing the metrics defined in [25]. Cells that could not be unambiguously assigned during annotation were excluded from the integration benchmarking. Benchmarking was carried out using either the latent embedding or the corrected protein expression matrices from the individual methods across the three preprocessing options with batch and ground truth cell annotations provided. For computational efficacy, evaluation of integration methods was performed on a random subset of 10 000 cells per batch.

#### Evaluation of protein imputation performance

To benchmark CytoVI’s performance for the imputation of missing protein markers, we utilized flow cytometry data of one individual PBMC donor from the *Nunez et al*. dataset [14]. The flow cytometry data was randomly down-sampled to 50 000 cells. CXCR3 and PD-1 were excluded from the analysis due to unspecific staining patterns. The resulting data was split into semi-synthetic batches by randomly assigning cells into one of two batch categories. Next we iteratively masked one marker at the time in one of the batches, trained a CytoVI model to integrate the two semi-synthetic batches and imputed the expression of the masked protein. To obtain robust estimates, we sampled each cell 50 times from the approximate posterior distribution and decoded its protein expression for the masked marker. The mean imputed expression across these 50 posterior samples was then compared to the observed protein expression for the respective cell. This procedure was repeated iteratively until all protein markers had been predicted. To estimate the uncertainty in the imputed protein expression we computed the standard deviation of the decoded protein expression across the 50 samples and assessed the relationship with the prediction error (the absolute difference between generated and observed protein expression) for the 50 samples across both, the cell and the protein axes. The imputation performance of CytoVI has been compared to cyCombine and KNN imputation, albeit the former one has been proposed to be used solely for visualization purposes due to the density sampling, which may hamper direct comparison [23]. KNN imputation has been performed using the scikit-learn implementation with 10 nearest neighbors. For cyCombine, we followed the standard workflow, using default parameters and z-score scaling of the arcsinh-transformed protein expression to generate the SOM labels. Imputation performance of the individual methods for each protein marker was compared by computing the Person’s and Spearman’s correlation coefficient between imputed expression and the held out observed expression. For markers that showed a clear bimodal expression pattern, the imputation performance was evaluated based on the binary marker positivity per cell. For this we fitted a two-component Gaussian Mixture model with the protein expression of an individual marker separately for each model and computed binary classification metrices between the observed ground truth marker positivity and marker positivity of each imputation method using scikit-learn.

### Imputation of missing markers to increase the dimensionality of cytometry assays

#### Imputation of missing proteins from two distinct flow cytometry antibody panels

To showcase CytoVI’s performance in integrating flow cytometry data from two distinct studies utilizing different antibody panels, we integrated data of PBMCs obtained from *Nunez et al*. [14] with flow cytometry data of PBMCs of an additional donor from a recent study investigating the systemic immune compartment of SARS-COV2 patients [27]. We used a weakly supervised approach by incorporating the cell type labels to inform the Mixture of Gaussian prior of the latent space *z*. The additional dataset from *Kreutmair et al*. interrogated 30 surface protein markers and was generated using a full spectrum cytometer [27]. The flow cytometry data of the *Kreutmair et al*. study was processed as described above for the other datasets. In short, the dataset was down-sampled to 100 000 cells, fluorescent intensities were transformed using a hyperbolic arcsin transformation employing a constant cofactor of 2000 and feature-wise scaled to a range of [0, 1]. The scaled protein expression was used to construct a nearest neighbor graph, compute UMAP and perform Leiden clustering using Scanpy. The resulting clusters were annotated into the major leukocyte subsets in the peripheral blood and harmonized to the cell type annotation in the *Nunez et al*. dataset. CD1c expression in the *Kreutmair et al*. data showed high unspecific staining patterns and was excluded from the integration. The concatenated dataset of the two studies was characterized by a shared backbone set of the following overlapping protein markers: CCR4, CCR7, CD123, CD14, CD16, CD19, CD27, CD3, CD38, CD4, CD45RA, CD56, CD8, CD95, HLA-DR. A CytoVI model was trained using default parameters controlling for the dataset identifier as batch covariate and incorporating the harmonized cell type annotations to weakly inform the prior of the latent space. Similar as before, a neighbor graph was constructed from the joint latent space and used for visualization using UMAP. Missing markers were imputed by sampling ten times from the approximate posterior distribution, generating the decoded protein expression for all markers and computing the mean across the sample dimension.

#### Generation of an integrated B cell maturation altas across 350 markers

For the generation of the imputed single cell B cell maturation atlas, mass cytometry data from *Glass et all*. have been accessed [31]. This dataset comprised PBMCs of two healthy donors each measured across 12 distinct mass cytometry antibody panels resulting in 24 unique samples. Single viable B cells were obtained from the mass cytometry.fcs files using manual gating in FlowJo and imported into Python. As in the original publication CD138 was excluded from the analysis due to unspecific staining. The mass cytometry data of the individual samples was transformed using a hyperbolic arcsin transformation with a constant cofactor of 5 and feature-wise scaled to a [0, 1] range. Due to the computationally expensive downstream analysis of trajectory inference methods, the dataset was down-sampled to 48 000 B cells in equal proportions from each sample [33]. A CytoVI model was trained by controlling for the sample key and only encoding the relevant B cell markers that were shared across the antibody panels: CD24, CD27, CD38, IgD, IgM. As the B cell population was expected to comprise a relatively homogenous population, we assumed an isotropic prior for the latent space of size *d* = 4. The trained CytoVI model was utilized to impute the non-overlapping protein markers by sampling ten times from the approximate posterior distribution, decoding the cells through the generative network to obtain the protein expression samples and computing the mean across the sample axis. Diffusion maps were generated by first computing the top 50 principal components, constructing a nearest neighbors graph based on this representation, and subsequently deriving the diffusion components using Scanpy [32]. Leiden clustering was applied to the nearest neighbor graph in conjunction with manual annotation in order to dissect the B cell maturation atlas into the major B cell subsets. Pseudotime of the B cell trajectory was estimated using Palantir [35], employing 5 components and otherwise default parameters, and using diffusion pseudotime as implemented in Scanpy (default parameters) [34]. Cell fate modeling was performed using Cell-Rank’s pseudotime kernel on a random subset of 20 000 cells of the integrated B cell maturation atlas [33]. Since Palantir and diffusion pseudotime showed highly similar results, the diffusion pseudotime was utilized to construct the pseudotime kernel in conjunction with the Generalized Perron Cluster Cluster Analysis (GPCCA) estimator. Terminal and initial states were set using prior knowledge about the underlying B cell biology. Fate-probabilities were computed using default parameters and the top 50 protein markers were identified by correlating the individual protein expression with the probabilities of B cells to differentiate towards the non-class switched memory B cell fate. Visualization of the protein expression dynamics in CellRank has been carried out by fitting a Generalized Additive Model with a Gaussian distribution weighting each cell’s contribution to each trajectory according to its fate probabilities. Differential protein expression between class-switched and non-class switched memory B cells was performed using CytoVI’s generative model **(see model Methods)**, while controlling for batch variation.

### Cross-technology integration and modality imputation

#### Imputation of morphological features from mass cytometry data

To showcase the capability of CytoVI to integrate distinct antibody-based single cell technologies with each other, we utilized the annotated and preprocessed mass cytometry data of PBMCs from [15] and flow cytometry data of PBMCs from [14] as described above. The dataset shared a common backbone of the following protein markers: CCR4, CCR7, CD127, CD14, CD16, CD19, CD27, CD3, CD38, CD4, CD45RA, CD56, CD69, CD8, FOXP3, HLA-DR, TCR*γδ*. The cell type annotations between the two datasets were harmonized by assigning leukocyte subsets to the next coarse cell type annotation level and were used to weakly inform the prior of the combined latent space *z*. A CytoVI model was trained using default hyperparameters. As described before a nearest neighbor graph was computed based on the latent representation of the trained CytoVI model and utilized for UMAP using Scanpy. We generating 10 samples from the approximate posterior distribution, decoded the union of features and computed the mean across the sample axis in order to obtain the imputed expression of all markers measured across the two studies, including the morphological features FSC and SSC. The mean expression of the imputed FSC and SSC per leukocyte subset in the mass cytometry data was compared to the mean of the measured FSC and SSC in the flow cytometry data by computing the Pearson’s correlation coefficient.

#### Automated detection of T cell states associated with disease in BNHL patients

To automatically detect T cell states associated with disease using CytoVI, we accessed raw flow cytometry data from a recent study investigating the T cell landscape in lymph nodes of non-Hodgkin B cell lymphomas [40]. The dataset comprised 63 lymph nodes samples that were measured in four batches by two overlapping antibody panels (12 surface and intracellular protein markers per panel) using conventional flow cytometry. Raw flow cytometry data was subjected to manual gating in FlowJo in order to identify viable singlet T cells and eliminate fluorescent aggregates. The individual batches were imported into Python and individually transformed using a hyperbolic arcsin transformation with a constant cofactor of 500 and scaled to the range [0, 1]. The data of the individual batches were merged into a single AnnData object yielding a shared backbone of the following proteins between the two flow cytometry panels: CD45RA, CD3, FOXP3, CD69, CD8, Ki67, CD4, PD-1. The resulting dataset was down-sampled to 10 000 T cells per patient. A CytoVI model was trained using default hyperparameters controlling for the observed batch covariate. Visualization of the latent space and imputation of non-overlapping markers was performed as described above. In order to obtain a reference annotation with respect to the original study, Leiden clustering of the CytoVI latent space and manual annotation was performed to identify the same T cell subsets as proposed by the original study authors [40]. To automatically detect T cell states associated with the different disease entities in a label-free manner, we utilized CytoVI’s differential abundance module **(see model Methods)**. In short, we aggregated the approximate posterior distribution for each patient and computed the log-median ratio of patients within a given disease entity relative to all other patients. Thereby, we retrieved a differential abundance score for each cell and each disease entity that can be used to detect regions in latent space that are characteristic for the respective disease entity. To identify differentially abundant T cell clusters the differential abundance scores for each cell and disease entity were concatenated to the latent representation of each cell and subjected to k-means clustering using scikit-learn. The confusion matrix between the identified differential abundance clusters and the ground truth cell annotation from the original authors was computed in Scanpy and rearranged based on the linear sum assignment. To evaluate the consistency of differentially abundant T cell clusters between the two applied flow cytometry panels, we computed the intra-class correlation coefficient (ICC) for the relative frequencies of these clusters, using the pingouin library.

#### Transcriptome imputation in flow cytometry data of BNHL patients

To impute the transcriptome of cells measured in the flow cytometry data of the BNHL cohort, we accessed paired CITE-seq data consisting of 70 oligonucleotide-tagged antibodies measured across 16 batches by *Roider et al*. [40]. To obtain batch corrected protein and RNA expression estimates, the 4000 most highly variable transcripts were selected and a TotalVI model was trained using default parameters [17]. Denoised and batch corrected protein and RNA expression was obtained by generating 10 samples from the approximate posterior distribution, generating the batch-corrected expression for each sample and computing the mean across the sample axis. Similarly as for the flow cytometry data, the resulting protein expression was transformed using a hyperbolic arcsin transformation using a constant cofactor of 5 and scaled to a range between [0, 1]. The protein expression of the CITE-seq dataset was merged with the flow cytometry data and standard-scaled. Cell type labels provided with the CITE-seq data were harmonized to the labels generated for the flow cytometry data. A CytoVI model was trained to integrate the flow cytometry and the protein expression data of the CITE-seq dataset using an adversarial loss scaled to 4 and a mixture of Gaussian prior that was weakly informed by the harmonized cell type labels. The integration performance was evaluated by comparing the annotation of each cell in the flow cytometry dataset to the annotation of its nearest neighbor in the CITE-seq data based on the integrated latent space and computing the confusion matrix between the labels. The joint latent representation was utilized by CytoVI’s imputation module to impute the transcriptome for cells measured by flow cytometry **(see model Methods)**. Imputation performance was benchmarked by comparing the batch-corrected protein expression in the flow cytometry dataset to its corresponding imputed transcript expression after log1p transformation and computing Spearman’s correlation coefficient. The protein-transcript relationship in the flow cytometry data was compared to the observed relationship in the CITE-seq data by correlating the denoised and scaled protein expression to the log1p transformed denoised expression of its corresponding transcript. Imputed transcripts were ranked using the log fold change for cells of the differential abundance cluster compared to all other T cells in the flow cytometry data using Scanpy and visualized in an MA plot. For the validation of the imputed transcript expression in the differentially abundant T cell cluster, a SCVI model was trained for the measured RNA data from the CITE-seq dataset using default parameters and correcting for the batch covariate [49]. The nearest neighbors graph was constructed based on the SCVI latent space and subjected to Leiden clustering using Scanpy in order to detect the cluster of *γδ* T cells and the corresponding gene expression was visualized in relation to the ground truth labels provided with the CITE-seq data.

### Automated classification of clinical flow cytometry data of CLL patients

#### Clinical characterization of CLL patients & ethical approval

The clinical analyses of initial diagnosis and disease course from CLL were obtained from patients treated at the University Hospital of Basel between 2020 and 2023 and the study was approved by the local ethic committee (BASEC 2024-00006). The median age of patients with detectable disease was 64 years, for MRD negative it was 62 years. Patient characteristics are summarized in **Supplementary Table S1**.

#### Flow cytometry data of clinical routine analysis

The clinical analyses of initial diagnosis and disease course in CLL patients were performed on blood samples using 10-color multiparameter flow cytometry (Lyrics, BD). The samples were analyzed sequentially for each patient as they were presented in the clinic, resulting in technical batches comprising one patient each. Collected samples were subjected to red blood cell lysis with BD FACS Lysing Solution. Following lysis, cells were washed with phosphate-buffered saline. For immunophenotypic characterization of CLL, a standardized flow cytometry panel was employed, including fluorochrome-conjugated monoclonal antibodies directed against CD45, CD19, CD20, CD5, CD200, CD10, *κ* and *λ* immunoglobulin light chains. All antibodies were purchased from BD, with the exception of the kappa and lambda antibodies, which were obtained from Dako **(Supplementary Table S2)**. Flow cytometry data acquisition was performed on a BD FACSLyric (BD Biosciences). Data were collected using FACS-Diva software, and fluorescence compensation was adjusted using single-stained controls for each fluorochrome. Fluorescence-minus-one controls were included to establish appropriate gating strategies. At least 50 000 total events were acquired per sample for standard immunophenotyping, while for minimal residual disease (MRD) detection, a minimum of 500 000 events was required to achieve a sensitivity of 10^−4^. MRD negativity was defined as *<*1 CLL cell detectable per 10 000 leukocytes (10^−4^) using manual gating by a trained hematologist. Clusters of *<*20 events were considered undetectable. Within the positive group CLL clones with a relative frequency *>*2% was defined as high tumor burden. Within the no detectable disease control, two non-CLL patients with only reactive changes in the blood and two non-CLL lymphoma patients have been included. In total 50 non-detectable disease control and 45 positive samples were chosen for downstream analysis using CytoVI **(Supplementary Table S1**.

#### Generation of a reference model of CLL patients

To generate a latent representation of leukocytes in the clinical flow cytometry data while accounting for technical variation between staining protocols and data acquisition sessions, we applied the following preprocessing steps. First, doublets, debris, and fluorescent aggregates were removed using FlowJo. The cleaned data were then imported into Python using the readfcs library for further analysis. Similarly as for the other datasets, the cleaned fluorescence intensity data were transformed using a hyperbolic arcsin transformation and scaled to a range of [0, 1]. To train and annotate a well-defined reference model ten CLL patients and ten non-detectable disease controls were selected. A CytoVI model was trained using this reference dataset by explicitly controlling for the sample key as the batch covariate. A latent space of size *d* = 5 was optimized, utilizing a *k* = 5 component mixture of Gaussian prior and encoding all protein markers except the immunoglobulin *κ λ* light chains into the latent space. The immunoglobulin light chains have not been passed to the encoder network to optimize a latent space that does not separate between Ig*κ*- or Ig*λ*-restricted CLL clones. The latent representation of the trained model has been used to construct a neighbors graph that has been clustered using Scanpys Leiden clustering implementation and manually annotated into the most abundant immune populations including the *CD*20^*dim*^ *CD*5^+^ CLL cells. Relative frequencies of each immune population among total cells per patient have been quantified using a Mann-Whitney U test. Relative frequencies of CLL cells have been compare to the ground truth frequency determined by hematologists using manual gating by computing Pearson’s correlation coefficient.

#### Automated patient classification using transfer learning

For the automated annotation of cells and consecutive classification of patient samples, we compiled the remaining dataset into the query dataset that was not used to train the model. For this, we loaded the query data into the trained reference model using the transfer learning approach introduced with scArches into scvi-tools [45, 10]. Passing the query data through the pretrained inference network yielded the latent representation of each cell without prior training. Cell type labels of cells in the query dataset were imputed using label transfer based on the 20 nearest neighbor in the CytoVI latent space **(see model Methods)**. Similarly as for the reference dataset, the relative cell type abundance in the query data was compared to the hematologists classification using a Mann-Whitney U test and frequency of imputed CLL cells per patient was compared to the ground truth annotation using manual gating by computing Pearson’s correlation coefficient. To estimate the immunoglobulin light chain restriction indicative of a monoclonal population, we computed the log ratio of the scaled Ig*κ* to Ig*λ* intensity of the CLL cells of each patient separately. For the differentiation of CLL patients (low and high tumor burden combined) compared to no-detectable disease controls, we trained a decision tree classifier of a depth of three using the relative frequency of the imputed CLL cell labels per patient and the log Ig*κ* to Ig*λ*ratio of the CLL cells as input features using the scikit-learn implementation. The model was evaluated using 5-fold cross-validation and the mean and standard deviation of the area under the ROC for the binary classification was reported.

## Supporting information

Supplementary_Table_S1_CLL_metadataSupplementary_Table_S2_Antibodies

Supplementary_Table_S2_Antibodies

## Data availability

Raw clinical flow cytometry data of CLL patients generated in this study are available upon request. Flow cytometry data of healthy PBMCs that served as batch normalization controls for the study by *Nunez et al*. has been provided by the authors [14]. Mass cytometry data of healthy PBMCs from *Ingelfinger et al*. has been provided by the authors [15]. CITE-seq data of PBMCs from *Hao et al*. has been accessed from the National Center for Biotechnology Information’s Gene Expression Omnibus under the accession number GSE164378 [16]. Flow cytometry data of PBMCs from *Kreutmair et al*. has been accessed at doi.org/10.17632/ffkvft27ds.2 [27]. Mass cytometry data of healthy B cells from the peripheral blood generated by *Glass et al*. are accessible at under the Flow Repository accession number FR-FCM-Z2MA. Raw unprocessed mass cytometry data were provided by the study authors [31]. Flow cytometry data of BNHL patients generated by *Roider et al*. were accessed at doi.org/10.6084/m9.figshare.24915633 [40]. CITE-seq data of BNHL patients from *Roider et al*. can be accessed at the European Genome-phenome Archive (EGA) under accession number EGAS50000000342. Cell annotations of the CITE-seq data have been provided by the original study authors.

## Code availability

Code to reproduce the analysis in this manuscript is available at https://github.com/YosefLab/cytovi-reproducibility. CytoVI is available as an open source software library integrated in https://scvi-tools.org. The reference implementation of CytoVI is available at https://github.com/YosefLab/cytovi-reference-implementation.

## Supplementary Figures

**Supplementary Figure S1.**
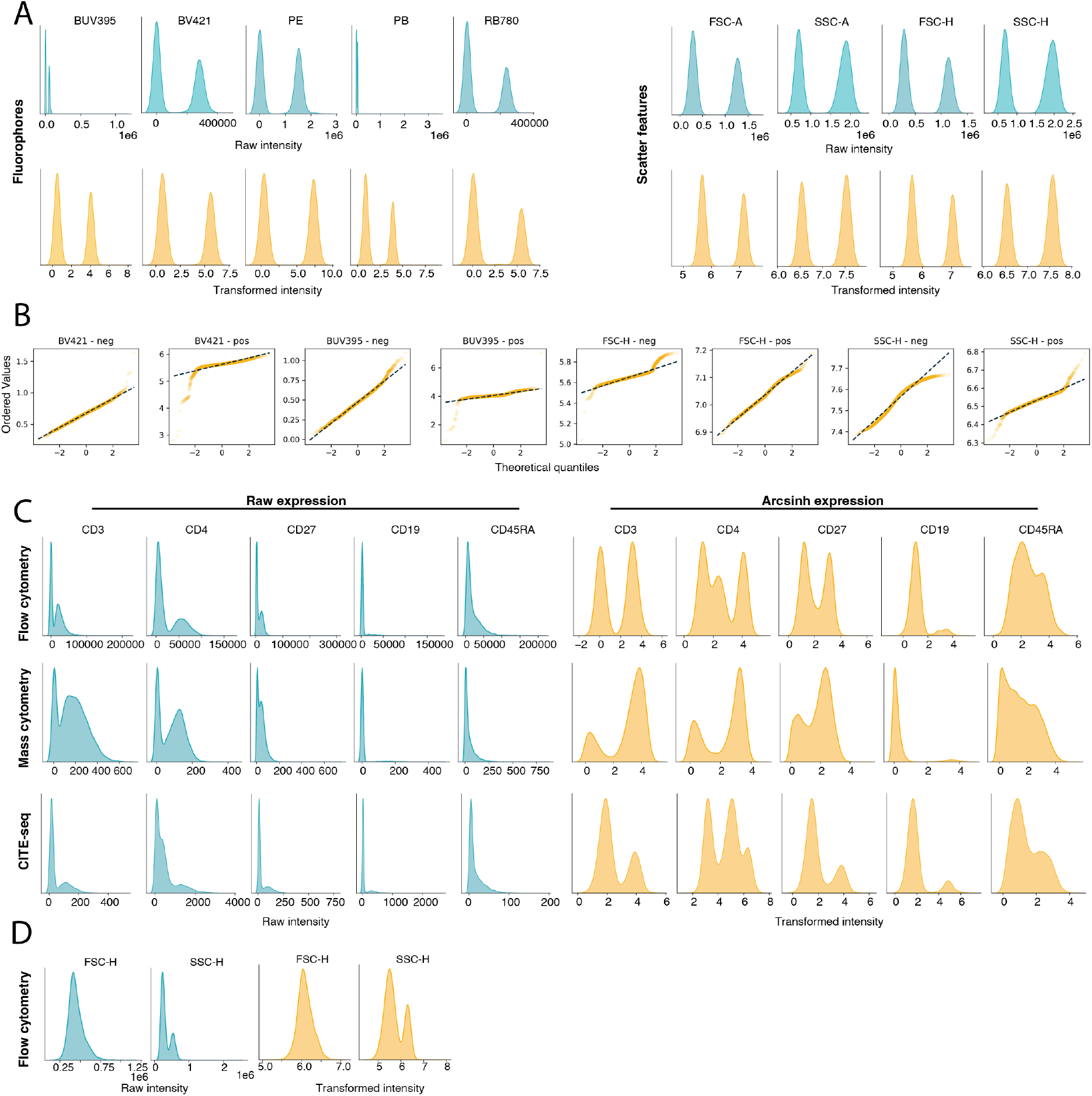
Transformed protein expression intensities across antibody-based single cell technologies can be approximated with a Gaussian distribution. **A:** Kernel-density estimate plots displaying the distributions of raw (blue) and hyperbolic arcsin transformed (yellow) fluorescence intensities (left panel) and scatter features (right panel) of single-stained microbeads (compensation beads) for the indicated fluorophores binding a homogeneous amount of antibodies. **B:** Quantile plots displaying the ordered values and the theoretical quantiles for the Gaussian distribution of the hyperbolic arcsin transformed fluorescence intensities of the positive (pos) and negative (neg) fraction of microbeads stained using the indicated fluorescent antibodies. The dashed line depicts the least-squares regression for the sample data. **C:** Kernel-density estimate plots displaying the distributions of raw (blue) and hyperbolic arcsin transformed (yellow) fluorescence intensities of the indicated protein markers in PBMCs measured by flow cytometry (*Nunez et al*.), mass cytometry (*Ingelfinger et al*.) and CITE-seq (*Hao et al*.). **D:** Kernel-density estimate plots displaying the distributions of raw (blue) and hyperbolic arcsin transformed (yellow) scatter features in PBMCs measured by flow cytometry. FSC = forward scatter, SSC = side scatter.

**Supplementary Figure S2.**
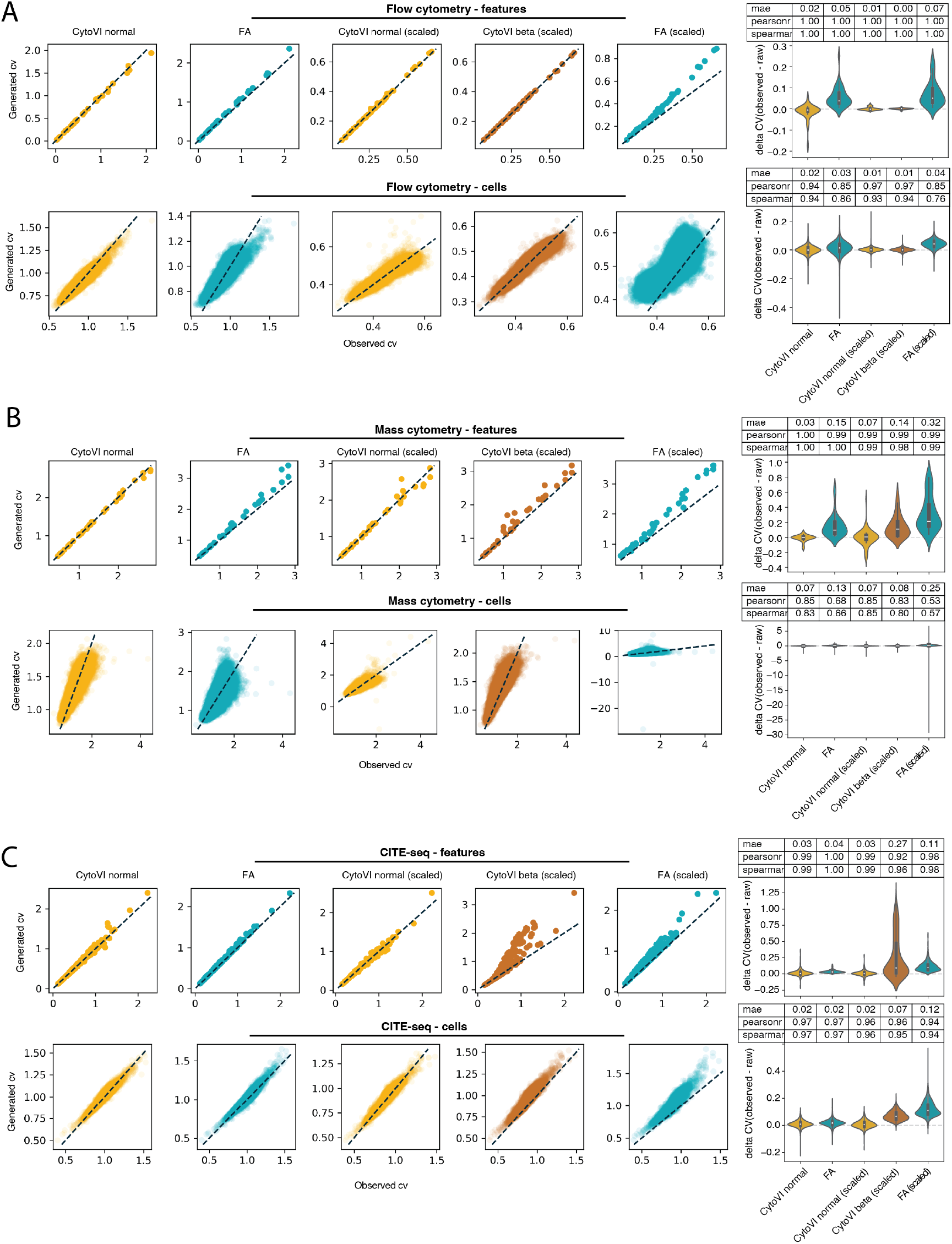
CytoVI generates protein expression estimates that closely match the observed data. **A, B, C:** Scatter plots (left panel) displaying the posterior predictive checks of the coefficient of variation (CV) across proteins (upper panel) and cells (lower panel) in the PBMC datasets measured by flow cytometry (**A**, *Nunez et al*.), mass cytometry (**B**, *Ingelfinger et al*.) or CITE-seq (**C**, *Hao et al*.). Posterior predictive checks were performed by computing the mean of the CV across posterior predictive samples obtained from CytoVI utilizing a Gaussian distribution with or without prior scaling of the data (CytoVI normal and CytoVI normal (scaled)), using a beta distribution (CytoVI beta (scaled) or using Factor Analysis (FA and FA (scaled)) as a baseline method. Violin plots (right panel) depict the prediction error of the CV for the indicated methods across all proteins (top panel) or all cells (bottom panel) with associated summary statistics displayed in the table above.

**Supplementary Figure S3.**
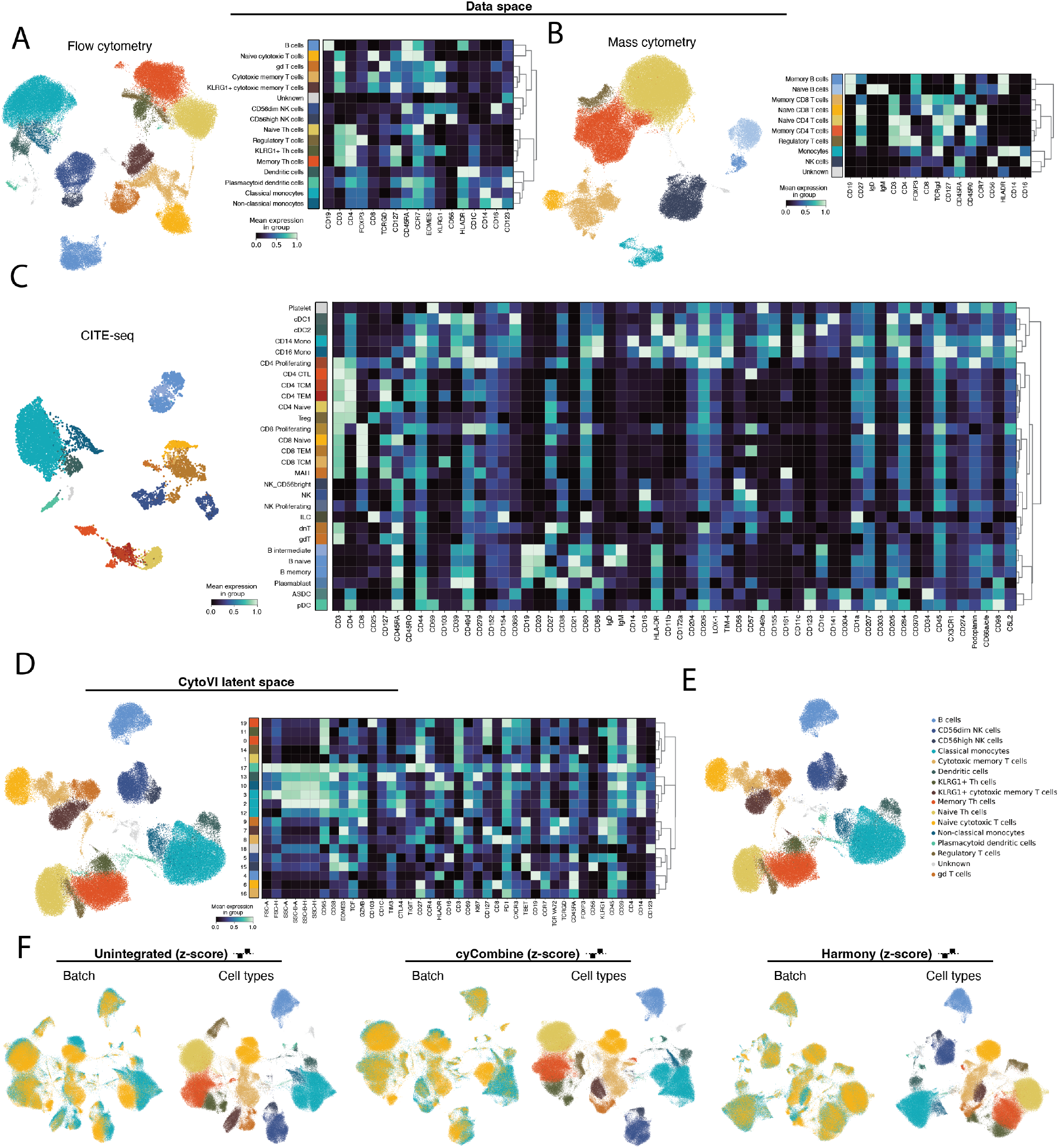
CytoVI controls for technical variation but preserves biological variation. **A, B, C:** UMAP (left panel) of the data space of PBMCs measured by flow cytometry (**A**, *Nunez et al*.), mass cytometry (**B**, *Ingelfinger et al*.), and CITE-seq (**C**, *Hao et al*.). Cells are colored by the ground truth cell type annotations that have been either provided with the original publication of the data **(C)** or were manually annotated **(A, B**. Heatmap (right panel) displaying the column-scaled mean protein expression intensities for the indicated immune populations. **D:** UMAP of the CytoVI latent space for PBMCs measured by flow cytometry (left panel). Cells are colored by the Leiden clusters obtained from the CytoVI latent space. Heatmap (right panel) displaying the column-scaled mean protein expression intensities for the indicated immune populations obtained from clustering the CytoVI latent space. **E:** UMAP of the CytoVI latent space for PBMCs measured by flow cytometry colored by their ground truth cell type label. **F:** UMAP of the unintegrated data space (left panels), the cyCombine corrected expression (middle panel) and the integrated embedding obtained from Harmony for two biological replicates of the flow cytometry samples measured in two different batches. Cells are colored by batch and the ground truth cell type annotation (displayed in **A**). In each of the three cases, the arcsinh-transformed data has been feature-wise z-score scaled prior to batch integration.

**Supplementary Figure S4.**
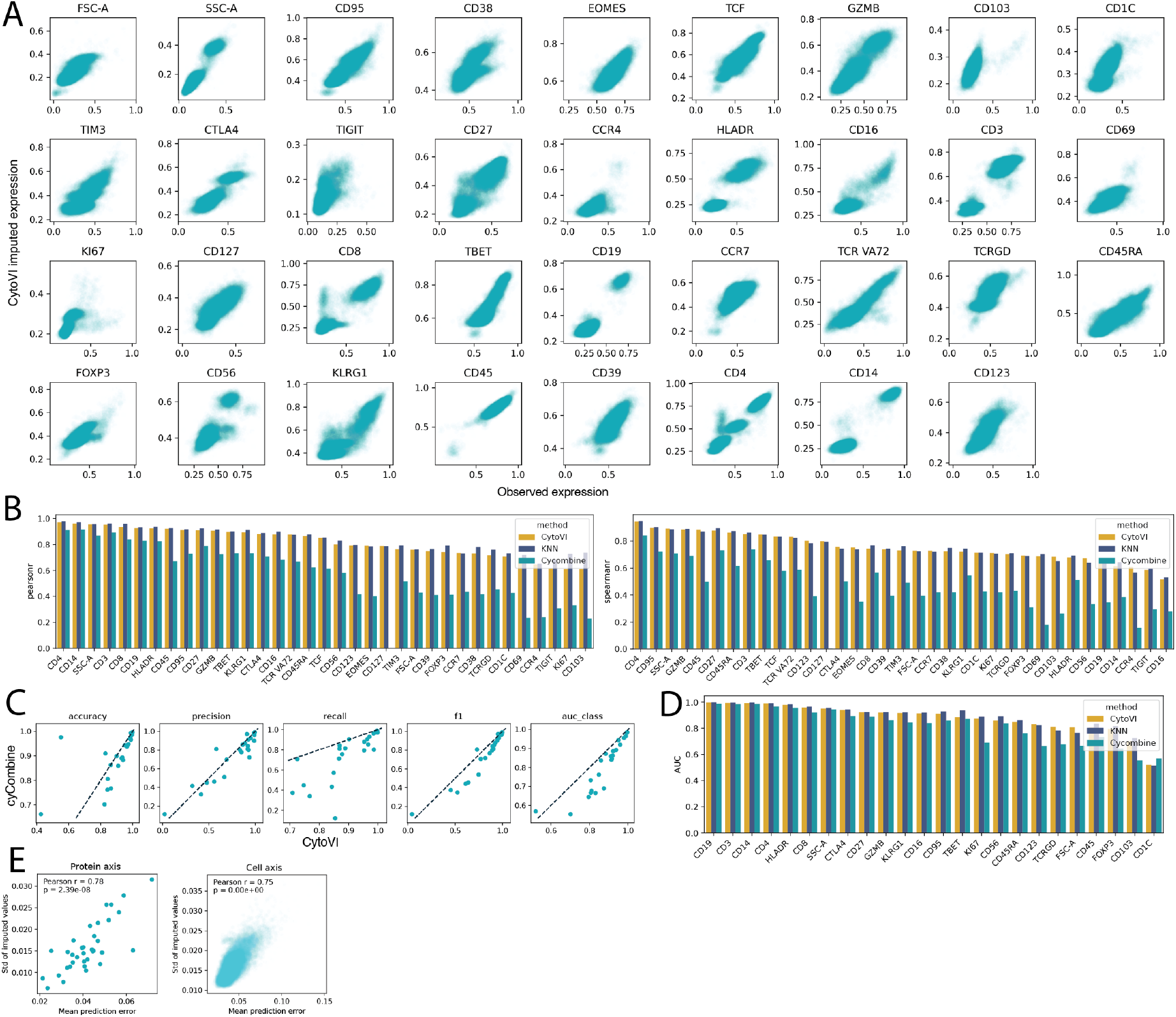
CytoVI accurately imputes protein expression in semi-synthetic experiments. **A:** Scatter plots displaying the observed and imputed protein expression for the indicated markers in the flow cytometry PBMC dataset. Cells of a single PBMC sample were randomly split into two batches, one marker at the time was masked in one of the batches, imputed using CytoVI and compared to the held-out protein expression. **B:** Bar plots displaying the Pearson’s (left panel) and Spearman’s (right panel) correlation coefficients between the observed and the imputed protein expression of the semi-synthetic imputation experiment using CytoVI, cyCombine and KNN. **C:** Scatter plots comparing the binary classification performance of cyCombine and CytoVI for imputed protein expression across different proteins. For proteins exhibiting a bimodal expression pattern, both observed and imputed protein expression values were classified into marker-positive and marker-negative classes using a two-component Gaussian mixture model fitted for each feature. The dashed line represents the identity line. **D:** Bar plots displaying the area under the ROC (AUC) for the binary classification of the indicated markers for each imputation method. **E:** Scatter plots showing the standard deviation (Std) of imputed protein expression across samples (left panel) and cells (right panel) along with the corresponding mean imputation error (compared to the observed expression) across samples using CytoVI. Samples were generated by sampling from the posterior distribution and passing the sample through the generative network.

**Supplementary Figure S5.**
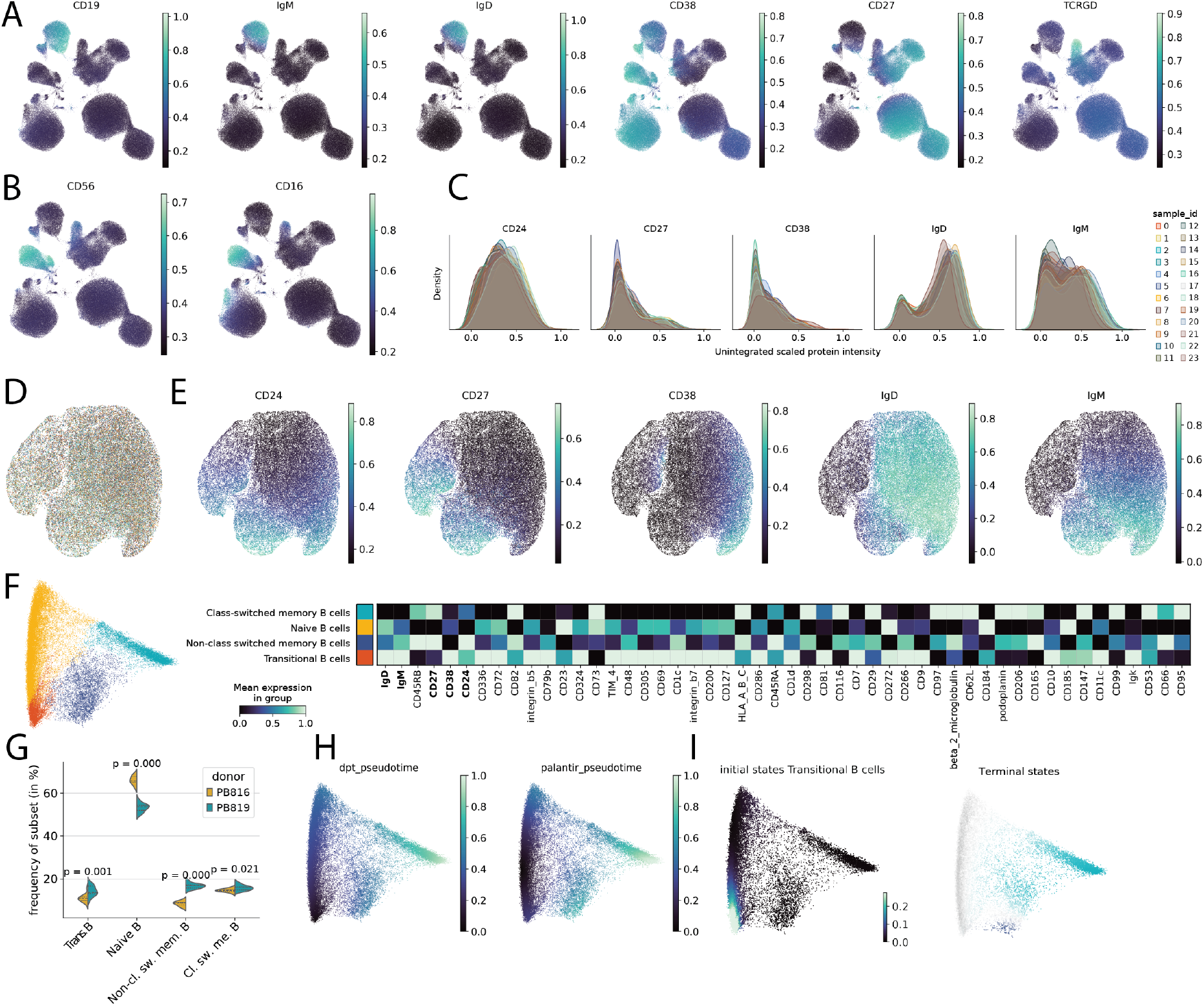
CytoVI integrates cytometry data from distinct antibody panels and imputes missing proteins. **A, B:** UMAP of the integrated CytoVI latent space of PBMCs from two distinct flow cytometry studies (*Nunez et al*., *Kreutmair et al*.) displaying the imputed expression of the indicated proteins. **C:** Kernel-density estimate plots displaying the uncorrected expression of the B cell markers in peripheral human B cells measured across 12 overlapping mass cytometry panels and two donors (resulting in 24 unique samples; *Glass et al*.). The displayed markers represent the backbone markers that are shared across the 12 antibody panels. **D:** UMAP of the CytoVI latent space of the B cell dataset colored by sample identifier. The color legend is displayed in **C. E:** UMAP of the CytoVI latent space of the B cell dataset displaying the batch-corrected expression of the backbone markers. **F:** Diffusion map (left panel) of the imputed protein expression profile of 350 surface markers of the B cell maturation atlas. Coloring indicates cell type annotations obtained from clustering the imputed data space followed by manual annotation. Heatmap (right panel) displaying the column-normalized mean imputed protein expression profiles of the 50 most variable markers between the indicated B cell subsets. Backbone markers that were measured in all of the antibody panels are highlighted in bold. **G:** Violin plots displaying the frequency of indicated B cell subsets for the two healthy donors measured across the different antibody panels (N = 12). P values correspond to a paired Wilcoxon test for the indicated B cell populations between the donors. **H:** Diffusion maps of the integrated B cell maturation atlas displaying diffusion pseudotime and Palantir pseudotime. **I:** Diffusion maps displaying the initial and terminal states of the B cell trajectory determined using CellRank.

**Supplementary Figure S6.**
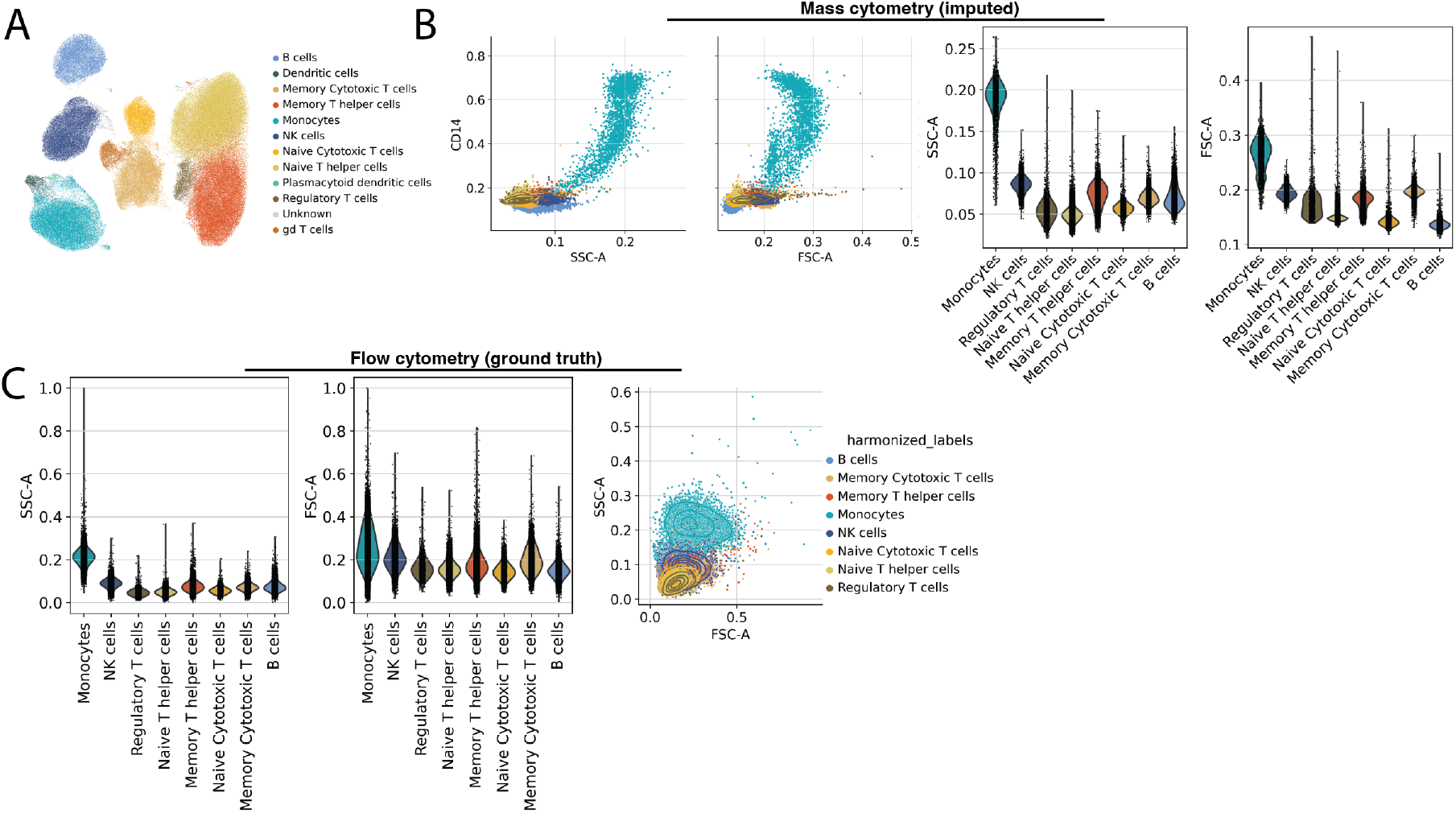
CytoVI imputes morphological features in mass cytometry data. **A:** UMAP of the CytoVI latent space displaying the integration of PBMCs measured by mass (*Ingelfinger et al*.) and flow cytometry data (*Nunez et al*.). Cells are colored by the harmonized cell labels between the datasets obtained from the ground truth annotation as defined in **Supplementary Figure S3A and B. B:** Scatter plots displaying CD14 in and the imputed scatter features in the mass cytometry data (left panel). Cells are colored by the harmonized cell labels between the datasets. Violin plots displaying the imputed scatter features for the indicated immune populations in the mass cytometry dataset (right panel). **C:** Violin plots displaying the measured ground truth scatter features for the indicated immune populations in the flow cytometry dataset (left panel). Scatter plots displaying the measured ground truth scatter features of the flow cytometry data (right panel). Cells are colored by the harmonized cell labels between the datasets.

**Supplementary Figure S7.**
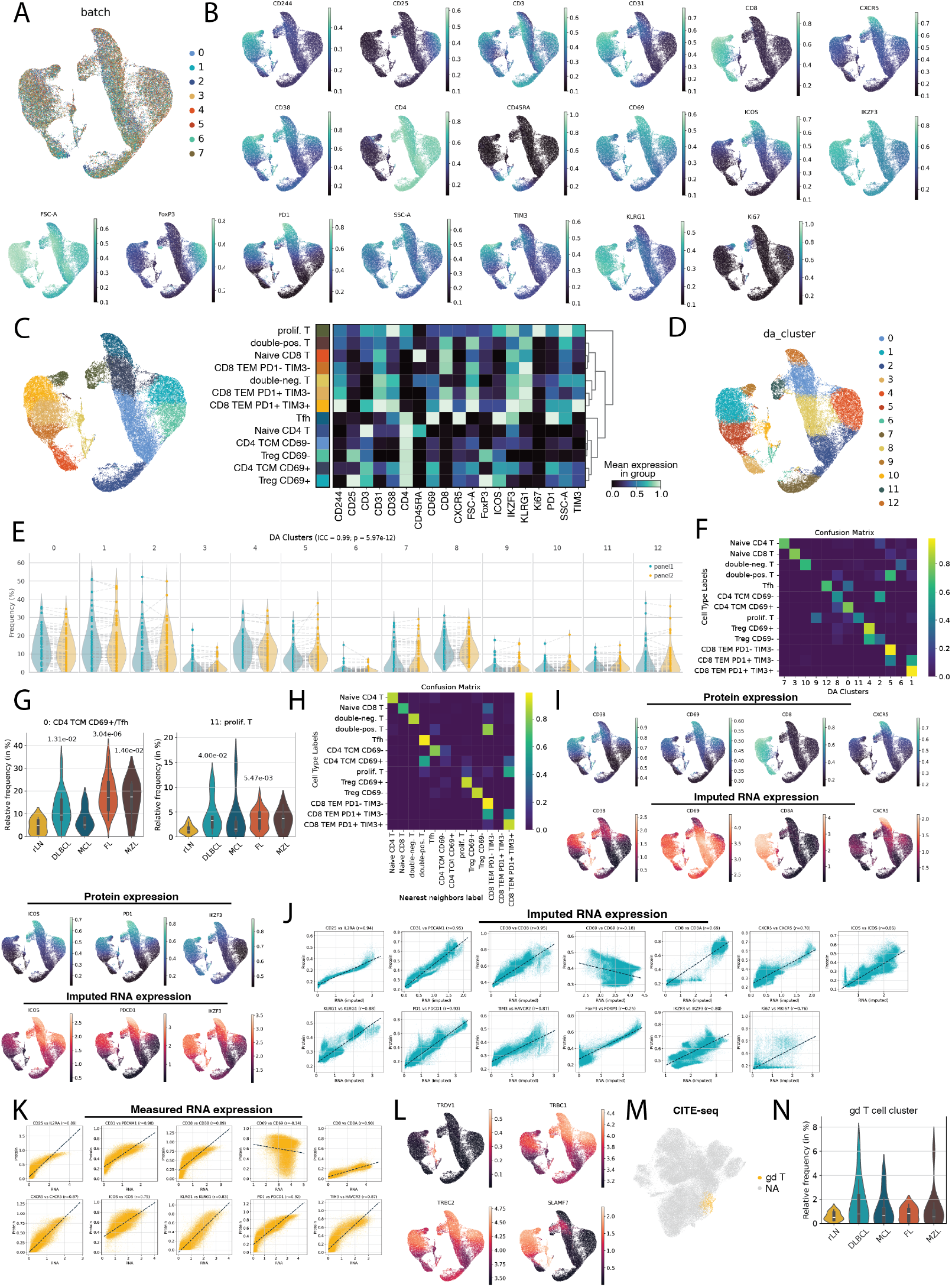
CytoVI automatically detects T cell states associated with disease in B cell non-Hodgkin lymphoma patients patients. **A:** UMAP of the CytoVI latent space of T cells obtained from lymph node samples of B cell non-Hodgkin lymphoma (BNHL) patients measured by flow cytometry (*Roider et al*.) colored by the batch in which the data was generated. **B:** UMAP of the CytoVI latent space of lymph node samples of BNHL patients displaying the imputed expression of the indicated protein markers. **C:** UMAP of the CytoVI latent space of T cells of BNHL patients colored by the cell state annotation analogous to *Roider et al*. (left panel). Heatmap displaying the column-scaled mean imputed expression profile of the indicated T cells states (right panel). **D:** UMAP of the CytoVI latent space of T cells of BNHL patients colored by the differential abundance (DA) clusters obtained from Leiden clustering of the per-entity differential abundance scores concatenated to the CytoVI latent space. **E:** Violin plots displaying the relative frequency of the differential abundance T cell clusters per sample separated by the panel that was used to measure the cells. The dashed line indicates samples that correspond to the same patient but where measured using a different antibody panel. Statistics indicate the intra-class correlation coefficient (ICC) across all clusters and the associated p value. **F:** Heatmap displaying the confusion matrix between the cell state annotations analogous to *Roider et al*. and the differential abundance clusters. **G:** Violin plots displaying the relative frequencies of the indicated differential abundance clusters across the different disease entities. rLN = resected lymph node (N = 7), DLBCL = diffuse large B cell lymphoma (N = 17), MCL = Mantel cell lymphoma (N = 9), MZL = marginal-zone lymphoma (N = 6), FL = follicular lymphoma (N = 24). P values refer to pair-wise Mann-Whitney U tests in relation to resected control lymph nodes (rLN). **H:** Heatmap displaying the confusion matrix between the cell state labels of the flow cytometry data of BNHL patients and the nearest neighbor annotation in the integrated CytoVI latent space for each cell in the CITE-seq data from *Roider et al*. To estimate the nearest neighbor labels a CytoVI model was trained on both technologies. **I:** UMAP of the CytoVI latent space of flow cytometry data displaying the denoised protein expression (top row) and the imputed gene expression of the corresponding transcript (bottom row) for the indicated markers. **J:** Scatter plots displaying the relationship between denoised protein expression in the flow cytometry data and the imputed gene expression of its corresponding transcript. Dashed line indicates the linear regression fit. The indicated correlation coefficient refers to Spearman’s correlation coefficient. **K:** Scatter plots displaying the relationship between denoised protein expression in the CITE-seq data and the denoised gene expression of its corresponding transcript using TotalVI. Dashed line indicates the linear regression fit. The indicated correlation coefficient refers to Spearman’s correlation coefficient. **L:** UMAP of the CytoVI latent space of flow cytometry data showing the imputed gene expression of the indicated transcripts. **M:** UMAP of the SCVI latent space of the transcriptome of the CITE-seq dataset highlighting the *γδ*T cell cluster obtained from Leiden clustering of the SCVI latent space. **N:** Violin plots displaying the relative frequency of the *γδ*T cell cluster obtained from Leiden clustering of the SCVI latent space among each patient in relation to the tumor entity (rLN: N = 8; DLBCL: N = 12; MZL: N = 12; FL: N = 11; MCL: N = 8).

**Supplementary Figure S8.**
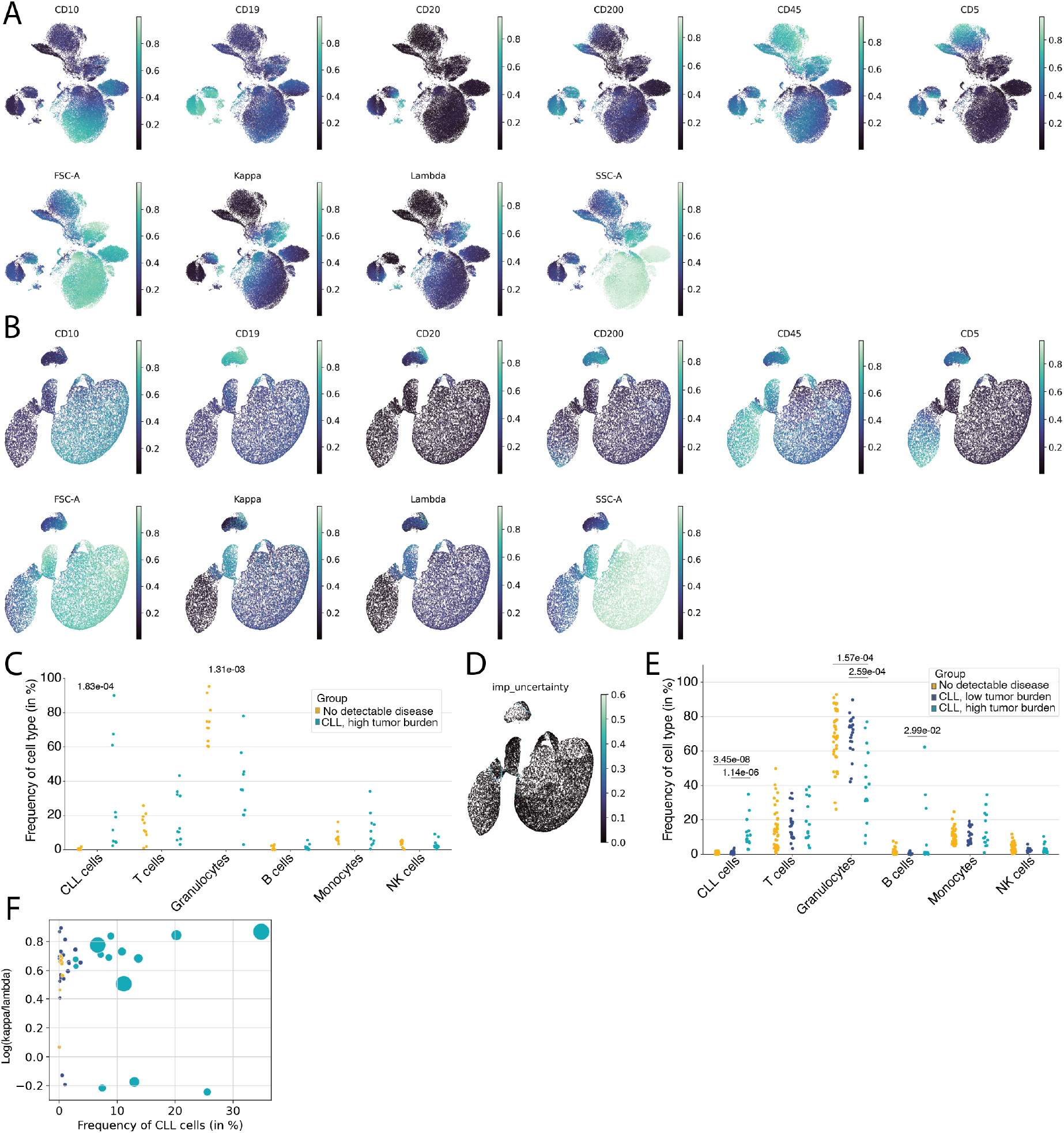
CytoVI can be applied to automate the diagnosis of chronic lymphocytic leukemia patients via transfer learning. **A, B:** UMAP of the unintegrated data space **(A)** and CytoVI latent space **(B)** of flow cytometry data of ten chronic lymphocytic leukemia (CLL) patients and ten no-detectable disease controls (comprising the reference dataset) displaying the arcsinh transformed protein expression of the indicated markers. **C:** Strip plots showing the relative frequencies of the indicated immune populations of high tumor burden CLL patients (teal; N = 10) and no-detectable disease controls (yellow; N = 10) in the reference dataset. P values refer to the two-sided Mann-Whitney U test. **D:** UMAP of the CytoVI latent space of the query dataset obtained via transfer learning displaying the uncertainty of the label transfer. The uncertainty corresponds to the relative frequency of the most abundant label among the nearest neighbors in the reference dataset for each cell in the query dataset based on the CytoVI latent space. **E:** Strip plots displaying the relative frequencies of the automatically-detected immune populations of high tumor burden CLL patients (teal; N = 14), low tumor burden CLL patients (blue; N = 21) and no-detectable disease controls (yellow; N = 40) in the query dataset. P values refer to the two-sided Mann-Whitney U test. **F:** Scatter plot showing the log(Ig*κ*/Ig*λ*) of the automatically-detected CLL cells in relation to the relative frequency of the CLL cells for each sample in the query dataset. Color corresponds to the diagnostic groups displayed in **E**. Dot size corresponds to the ground truth relative frequency of CLL cells determined by expert manual gating.

## Supplementary Tables

**Supplementary Table S1: Clinical characteristics of the CLL cohort**. Table indicating the diagnosis for each patient utilized in the CLL cohort, the presence of the Matutes criteria in the clonal population, the ground truth relative frequency of CLL cells (among B cells and total leukocytes) assessed using manual expert gating and the light chain restriction of the CLL clones.

**Supplementary Table S2: Antibodies used for flow cytometry analysis of CLL patients**. Table displaying the antibody panel including fluorophores, clones and manufacturers for antibodies used to analyze the CLL cohort.

## Acknowledgments

We would like to thank David Glass for providing the raw data of the B cell maturation atlas, Nico Nunez for providing the raw data of the normalization controls in the flow cytometry study and Donnacha Fitzgerald, Hosna Baniadam, Sascha Dietrich and Wolfgang Huber for providing the raw data for the BNHL CITE-seq data. We thank Pascal Schlaepfer for support with anonymization of the clinical flow cytometry data. We thank Tomer-Meir Salame, Florian Mair, Adam Gayoso, Burkhard Becher and Bertram Bengsch for the critical discussions. I.A. is an Eden and Steven Romick Professorial Chair, supported by the HHMI International Scholar Award, funded by the European Union ERC advanced grant (no. 101055341-TROJAN-Cell), European Union (IMMEDIATE EU#: 101095540), the MBZUAI/WIS joint program on artificial intelligence, the Deutsche Forschungsgemeinschaft (DFG, German Research Foundation) – Project-ID 259373024 – TRR 167, and the Israel Science Foundation grant no. 1944/22, co-funded by the European Union (ERC, MiTE, 101123436). This research was supported by a Research Professorship grant from the Israel Cancer Research Fund, United States-lsrael Binational Science Foundation (BSF), Ministry of Innovation, Science & Technology grant (1001703362(, Dwek Institute for Cancer Therapy Research, Moross Integrated Cancer Center, EKARD Institute for Cancer Diagnosis Research, Morris Kahn Institute for Human Immunology, Swiss Society Institute for Cancer Prevention Research, Elsie and Marvin Dekelboum Family Foundation, Lotte and John Hecht Memorial Foundation and the Schwartz Reisman Collaborative Science Program, Canada BrainResearch Fund (CBRF), Dr. Daniel C. Andreae, and the Larry and Judy Tanenbaum Family Foundation. This study was supported by the Center for Immunotherapy at the Weizmann Institute of Science. The work on this project was supported by the Chan Zuckerberg Initiative Essential Open Source Software grants EOSS4-0000000121 and EOSS6-0000000388 (N.Y.). It was also supported by the Chan Zuckerberg Initiative Silicon Valley Community Foundation - Single-Cell Biology Data Insights program (DAF2022-249318) and by NIH/NIDA award R61DA048444 (N.Y.). F.I. was a recipient of the Koshland price and was funded by an EMBO postdoctoral fellowship (ALTF 723-2022) and a MSCA postdoctoral fellowship (101106452; STIC-GBM). F.I. received funding from the Deutsche Forschungsgemeinschaft (DFG) research grant number 441891347 (SFB1479-OncoEscape).

